# Distinct internal states interact to shape food choice by modulating sensorimotor processing at global and local scales

**DOI:** 10.1101/2021.05.27.445920

**Authors:** Daniel Münch, Dennis Goldschmidt, Carlos Ribeiro

## Abstract

When deciding what to eat, animals evaluate sensory information about food quality alongside multiple ongoing internal states. How internal states interact to alter sensorimotor processing and shape decisions such as food choice remains poorly understood. Here, we use pan-neuronal volumetric activity imaging in the *Drosophila* brain to investigate the neuronal basis of internal state dependent nutrient appetites. We created a functional atlas of the ventral fly brain and find that metabolic state shapes sensorimotor processing across large sections of the neuropil. Reproductive state, in contrast, acts locally to define how sensory information is translated into feeding motor output. Thereby, these two states synergistically modulate protein-specific food intake and thus food choice. Finally, using a novel computational strategy, we identify driver lines innervating state-modulated regions and show that the newly identified borboleta region is sufficient to direct food choice towards protein-rich food. We therefore identify a generalizable principle by which distinct internal states are integrated to shape decision-making and propose a strategy to uncover and functionally validate how internal states shape behavior.

## Introduction

Behavior and decision-making depend on the ability to integrate sensory information with internal states. Depending on the current or future needs of an animal, the same sensory information can be evaluated differently and lead to distinct behavioral outcomes (Burgess et al., 2018; Corrales-Carvajal et al., 2016; Deutsch et al., 1989; Filosa et al., 2016; Griffioen-Roose et al., 2012; Lewis et al., 2015; Ribeiro and Dickson, 2010; Richter and Barelare, 1938; Root et al., 2011; Simpson and Abisgold, 1985; Steck et al., 2018; Vogt et al., 2021; Walker et al., 2015). Internal states such as hunger and thirst have long been known to change how sensory information guides behavioral choices (Burton et al., 1976). How internal states modulate sensory information across different levels of processing and how this changes decision-making is not clear. Furthermore, internal states alter behavior and decision-making in remarkably specific ways (Münch et al., 2020). This is thought to be driven by modality-specific alterations of sensory information processing (Rolls, 2015; Rolls et al., 1981; Steck et al., 2018). But if such sensory specificity extends across the sensorimotor arc remains to be characterized. Finally, while most studies focus on one internal state variable, in reality, animals are exposed to multiple, sometimes conflicting, internal needs. Despite being a fundamental question in neuroscience, we do not understand how the brain reads out and integrates information from multiple internal states to reach one specific decision.

Nutrients are key determinants of the health and well-being of animals (Simpson and Raubenheimer, 2012). Both under- and over-ingestion of specific nutrients can have severely deleterious effects on fitness (Carvalho-Santos et al., 2020; Levine et al., 2014; Margetts, 2004; Piper et al., 2017; Solon-Biet et al., 2014). As such, animals have evolved the ability to avoid or ameliorate specific nutritional deficiencies by choosing to eat from foods that contain the required nutrients (Münch et al., 2020). To do so, they integrate information about current and future internal states with food sensory information, especially taste (Leitão-Gonçalves et al., 2017; Rolls, 2015; Simpson and Simpson, 1992; Steck et al., 2018; Walker et al., 2015). The resulting so-called nutrient-specific appetites are especially relevant for carbohydrates, protein, and salt, which have been demonstrated to exist in animals ranging from invertebrates to humans (Beauchamp et al., 1990; Carvalho-Santos et al., 2020; Deutsch et al., 1989; Griffioen-Roose et al., 2012; Hughes and Wood-Gush, 1971; Leitão-Gonçalves et al., 2017; Richter and Barelare, 1938; Simpson and Raubenheimer, 2012). Thus, the phenomenon of nutrient-specific appetites represents a conceptually powerful and biologically highly relevant paradigm to study how internal states shape sensorimotor processing and decision-making (Münch et al., 2020). Importantly, nutrient-specific appetites are not only determined by the lack of specific nutrients, but result from the integration of metabolic with other states, such as reproductive state (Ribeiro and Dickson, 2010; Walker et al., 2015). In contrast to metabolic state, which is thought to represent the current nutritional needs of the animal, reproductive state represents future nutritional needs and hence shapes decision-making in a feed-forward, predictive way (Walker et al., 2017). Intriguingly, while evidence exists that metabolic state shapes food choice by altering sensory processing, reproductive state seems to modulate nutrient appetites by a different mechanism (Steck et al., 2018). The neuronal mechanisms by which mating affects nutrient-specific food choice and how and where along the sensorimotor arc reproductive and metabolic states are integrated to synergistically enhance protein intake remains to be understood.

While primary sensory neurons usually target well segregated brain regions, the downstream processing of sensory information is highly distributed, involving many classes of neurons at different computational levels (Allen et al., 2017; Pacheco et al., 2021; Portugues et al., 2014). Likewise, while specific regions like the prefrontal cortex have been linked to decision-making, especially value-based decision-making, newer data supports models in which decision-making involves a distributed network of many cortical and subcortical regions (Steinmetz et al., 2019). We lack an understanding of how these distributed dynamics are altered by different internal states and how these are functionally linked to decision-making.

Pan-neuronal imaging across large brain volumes at cellular resolution in small model-animals, as well as high-density electrophysiology recordings across the brain in rodents, have started to shed light on the distributed nature of neuronal computations underlying behavior and cognition (Urai et al., 2021). Pan-neuronal imaging techniques have been largely pioneered in larval zebrafish (Ahrens et al., 2012, 2013; Portugues et al., 2014) and have more recently been applied to other animals including *Drosophila*, *via* recordings at either cell-bodies or the neuropil (Aimon et al., 2019; Harris et al., 2015; Mann et al., 2017; Pacheco et al., 2021; Schrödel et al., 2013). While these studies have highlighted the distributed nature of sensory processing (Pacheco et al., 2021; Portugues et al., 2014) and the engagement of large parts of the brain in behaving animals (Aimon et al., 2019; Allen et al., 2017; Marques et al., 2019; Schrödel et al., 2013; Steinmetz et al., 2019), relatively little is known about the impact of internal states on sensorimotor processing. Emerging data suggest that internal states change brain activity at a more global scale (Allen et al., 2019), but the causal relationship of these changes with alterations in behavior remains unclear, as does the generalizability of this observation. The highly distributed nature of these signals emphasizes the importance of studying brain processes using more global recording approaches. However, such distributed recording approaches make it even more challenging to validate the functional relevance of the observed distributed neuronal changes.

Similarly to humans (Griffioen-Roose et al., 2012), *Drosophila melanogaster* also shows a switch in preference towards protein-rich foods when low on amino acids (Leitão-Gonçalves et al., 2017), a choice that relies to a large extent on gustatory information (Steck et al., 2018). In flies, taste neurons send their projections to the subesophageal zone (SEZ), an ancestral neuropil with similarities to the vertebrate brainstem that has been proposed to be key for mediating sensorimotor transformations in invertebrates (Gal and Libersat, 2010; Knebel et al., 2018; McKellar, 2016; Tastekin et al., 2018, 2015). Unlike most motor outputs that are generated in the ventral nerve cord, the proboscis motor neurons that drive the feeding motor program are also located in the SEZ (Gordon and Scott, 2009; McKellar et al., 2020; Schwarz et al., 2017). Housing both feeding-relevant input and output neurons, it is likely that most of the functionally relevant integration of internal state and food taste information takes place within the SEZ. This makes it theoretically possible to map the full sensorimotor loop and the sensorimotor transformations therein by studying this neuropil. A key challenge in doing so is that while both the taste neurons and the motor neurons mediating feeding have been studied in some detail, the largest part of the SEZ, including most of the circuitry downstream of the sensory neurons and the neurons controlling feeding, remains enigmatic.

To investigate how different internal states shape sensorimotor transformations and how this ensures appropriate food choice, we imaged neuronal activity across the SEZ of *Drosophila* using pan-neuronal volumetric activity imaging. We then compared responses to food taste stimuli across animals in different internal states *via* the generation of a functional atlas. We found that both internal states specifically altered sensorimotor processing of food containing the nutrient the animal needed. But while metabolic state acted across multiple layers of processing, mating exclusively increased activity in motor regions upon sensory stimulation. Importantly, the effect of reproductive state was only visible in protein-deprived females, providing a neuronal basis for the synergistic effect of mating on food choice. Using this functional SEZ atlas, we furthermore discovered novel SEZ regions that integrate sensory with metabolic state. In particular, a region we termed “borboleta” shows a strong and specific increase in responsiveness to protein-rich food upon protein deprivation. We then used a computational strategy to select candidate driver-lines with expression in this region. Thermogenetic activation experiments revealed that neurons innervating the borboleta region are sufficient to bias food choice towards yeast. Our work demonstrates two fundamentally different modes by which internal states act synergistically on the same feeding decision, and provides functional evidence that the borboleta region directs food choice toward proteinaceous food.

## Results

### *Drosophila* food choice is strongly modulated by multiple internal states

Animals adapt behavioral decisions to their needs in order to optimize fitness (Andermann and Lowell, 2017; Münch et al., 2020; Rangel et al., 2008). These needs are thought to be represented as internal states in the brain, altering how sensory information is processed to guide animals’ choices among different options. We have shown that when presented with the choice to feed on either a sucrose or a protein-rich yeast patch, *Drosophila melanogaster* adjusts its food choice to prioritize the intake of the nutrients it needs (Itskov and Ribeiro, 2013; Ribeiro and Dickson, 2010). When lacking dietary amino acids (AAs) due to protein deprivation, flies shift their preference towards the protein-rich food source, increasing yeast feeding and decreasing sugar feeding (Figure 1a). Furthermore, in AA-deficient females, mating further biases the preference of the female towards protein by increasing protein appetite without affecting sugar feeding ((Figure 1a). This dual state modulation is consistent with the principle that most decisions are not shaped by only one internal state, but by a plethora of continuously changing internal state variables. These have to be integrated by the brain to form appropriate behavioral adaptations. Accordingly, food choice is synergistically influenced by the metabolic state of the animal and its reproductive state. We therefore decided to use food choice as an ethologically relevant paradigm to dissect how internal states modulate sensorimotor processing and how different internal states are integrated to shape decisions.

**Figure 1.**
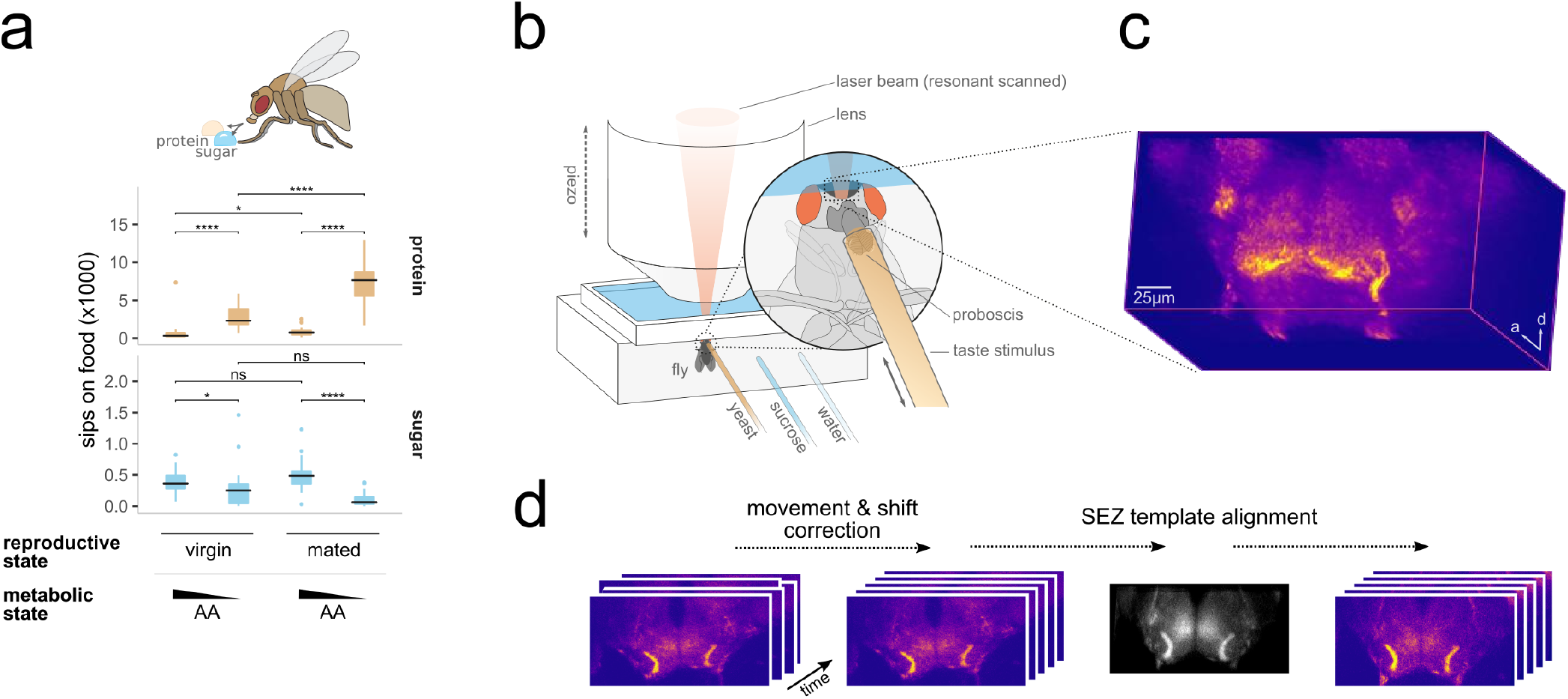
Internal states shape food choice and pan-neuronal volumetric imaging setup. **a**, Number of sips as measured by the flyPAD assay from 10 % yeast (protein, *beige*) and 20 mM sucrose (sugar, *blue*) by virgin or mated female flies of the genotype *57C10-GAL4,UAS-GCaMP6s*, fed for 10 days on a complete diet or a diet lacking protein. Boxplots indicate median, first, and third quartile, whiskers extend to the lowest and highest values that lie within 1.5 of the inter-quartile range of the box. Points represent data points outside of the whisker range. Holm corrected Wilcoxon rank-sum test, n = 22-28, ns P > 0.05, * P < 0.05, **** P < 0.0001. **b**, Schematic showing the imaging setup. A living fly is fixed to the recording chamber. A hole in the head capsule gives optical access to the brain. The proboscis is fixed in an extended position and stimulated with different taste solutions. **c**, An example volume rendering of a pan-neuronal volumetric SEZ recording. **d**, Schematic depicting the image preprocessing steps; imaging data are first corrected for movements within and shifts between recordings in 3D followed by an alignment to a common SEZ template to enable comparative analysis across animals in different internal states.

### Pan-neuronal volumetric imaging of brain activity in *Drosophila*

Functional imaging of specific neuronal populations has contributed significantly to our understanding of how neuronal circuits relay sensory information to shape behavior (Chiappe et al., 2010; Marella et al., 2006; Sachse et al., 2007). Yet, the neuronal computations underlying decision-making likely involve large numbers of neurons, distributed within and across neuropils (Urai et al., 2021). We therefore decided to tackle how internal states shape sensorimotor processing to inform decision-making by performing pan-neuronal calcium imaging in brains of flies in different internal states exposed to different gustatory food stimuli. We used a resonant scanning two-photon microscope in combination with a piezoelectric lens drive to volumetrically record food taste-induced neuronal activity in the brains of living flies, expressing the calcium reporter GCaMP6s pan-neuronally (Figure 1b). Our setup allows us to capture volumetric time series at a reasonably high resolution of 512 × 256 × 55 voxels with voxel dimensions of ~0.5 × 0.5 × 3.6 µm and a volume rate of ~ 1 Hz while stimulating the gustatory sensory neurons on the tip of the flies’ proboscis with different food solutions (Figure 1b and c; Supplementary Video 1). Flies in four different internal states (fully fed & virgin, fully fed & mated, low AAs & virgin, low AAs & mated) were stimulated twice with three different food stimuli (water, sucrose, and yeast) (Figure S1). After correcting for movement artifacts, volumetric time series were registered to a template brain to allow us to compare across flies and internal states (Figure 1d).

In the arthropod brain, similarly to the vertebrate brainstem, the subesophageal zone (SEZ) has the unique property of both receiving gustatory information and sending out motor commands that control food intake (McKellar, 2016; Scott, 2018) It is therefore reasonable to assume that the SEZ contains circuits involved in mediating the effects of internal states on feeding. We therefore decided to optimize the trade-off between imaging speed and resolution by focusing our pan-neuronal imaging on the ventral *Drosophila* brain, where the SEZ is located.

### A functional atlas of the *Drosophila* subesophageal zone

The SEZ comprises four ventral brain neuropils: the antennal mechanosensory and motor center, saddle, prow and the gnathal ganglion (GNG), with the GNG making up the largest part of the SEZ volume (Ito et al., 2014; Figure 2a). Most of what is known about the GNG is limited to the morphology of the projections of gustatory sensory neurons as well as the arborizations of feeding motor neurons (McKellar et al., 2020; Miyazaki and Ito, 2010; Schwarz et al., 2017; Thorne et al., 2004; Wang et al., 2004). This stands in contrast to other brain regions, for which there is a large body of work describing their functional and anatomical organization. Given the lack of a granular anatomical map of the SEZ, we aimed to apply an activity-based brain segmentation strategy to create a functional map of the SEZ. Such atlases serve as important tools to dissect activity across brain regions. To create the SEZ atlas, we used a group-level implementation of spatial independent component analysis (ICA; Varoquaux et al., 2010 to extract spatially independent components based on the activity observed in all the averaged 4D imaging stacks (Figure 2b). For each recording (water, sucrose, and yeast stimulation) of animals in one of the four different internal states (Figure S1), we calculated ΔF/F image stacks and averaged them across animals. We then ran the ICA algorithm across these data to extract spatial components (Figure S2a and b). Finally, we built a single binary SEZ atlas by applying thresholding, followed by erosion and dilation steps (see Materials and Methods for details). The final atlas consists of 81 largely bilaterally symmetric regions spanning the entire SEZ (Figure 2b, S3, and Supplementary Data 1). Our SEZ atlas comprises regions that correspond to most previously described sensory neuron projections (Figure 2c). These include, for example, the AMS1 and PMS4 gustatory neuron projection areas in the medial SEZ, as well as a set of horizontally layered regions in the anterior ventral SEZ that resemble the areas innervated by feeding motor neurons. Importantly, scrambling the temporal structure of the imaging data resulted in atlases with no obvious structure and no bilateral symmetry, validating the SEZ atlas we generated (Figure S2c and d).

**Figure 2.**
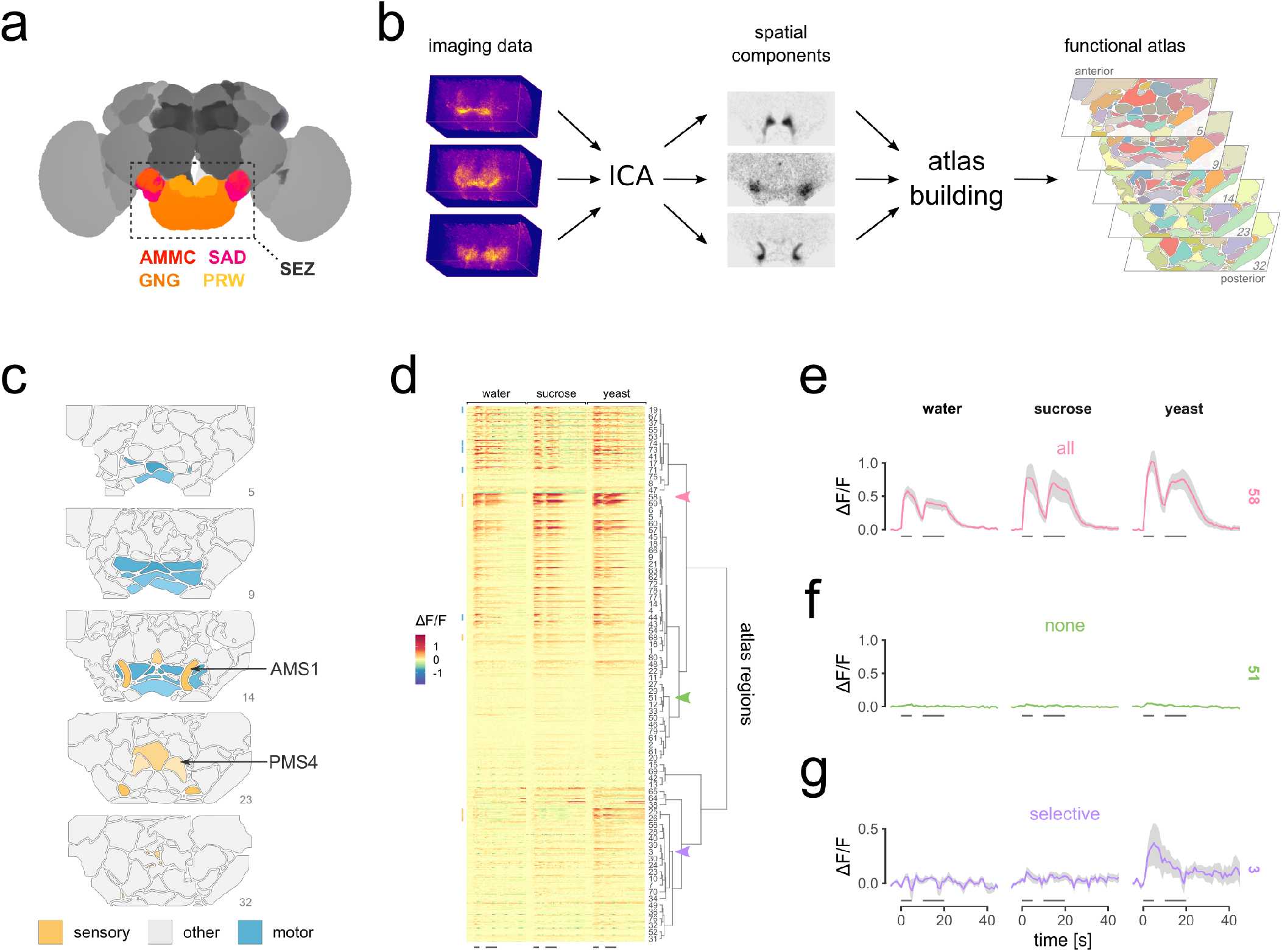
A functional atlas of the SEZ. **a**, Diagram of a fly brain with the SEZ and its major neuropils highlighted; anterior mechanosensory and motor center (AMMC), saddle (SAD), gnathal ganglion (GNG) and prow (PRW) in *magenta, red, orange* and *yellow* respectively. Segmentation from (Ito et al., 2014). **b**, Schematic depicting the process for creating the functional SEZ atlas. Spatial components are extracted from the imaging data using group-level ICA. The atlas is built by merging components into a single binary atlas using thresholding and dilation and erosion steps. In this and all future depictions of the atlas, numbers refer to the number of the slice being depicted. **c**, The final atlas contains 81 regions, some of which resemble described regions innervated by sensory projections- (*yellow*) and motor neuron arborizations (*blue*). **d**, Heatmap of activity time series extracted from each atlas region of individual mated, protein deprived animals stimulated with the three different sensory stimuli. The data is grouped by atlas region and within the region ordered by average activity strength. The order of atlas regions is based on hierarchical clustering of the average time series of each region across animals. Lines at the bottom depict the two consecutive stimuli. Arrowheads highlight regions for which traces are shown in (**e-g**). Example of a region showing response to all stimuli (**e**), to none of the stimuli (**f**), or responding selectively to some stimuli (**g**), respectively. Shaded areas in (**e-g**) indicate mean ± sem, *gray dashes* indicate stimulation, n = 8-9.

Next, we used the registered imaging data from each animal to extract voxel-averaged time series from each region of the atlas, which allowed us to generate comparable time series for each gustatory food stimulation across individuals (Figure 2d and S4). The resulting traces showed a wide variety of response patterns, with some regions responding to all stimuli (Figure 2e), some showing no response upon stimulation (Figure 2f), and some responding in a stimulus-specific manner (Figure 2g). Regions corresponding to the projection of bristle gustatory neurons showed generalized responses to gustatory stimuli, albeit to varying degrees (regions 58 & 59, PMS4) (Figure 2d and e). This is consistent with previous observations showing that all stimuli used here activate gustatory neurons projecting to PMS4 (Inoshita and Tanimura, 2006; Marella et al., 2006; Steck et al., 2018). In contrast, regions corresponding to the projection pattern of the yeast stimulus-specific peg gustatory neurons (regions 25 & 26, AMS1) showed largely yeast-specific sensory responses (Figure 2d) (Fischler et al., 2007; Sánchez-Alcañiz et al., 2018; Steck et al., 2018). Moreover, as previously suggested (Harris et al., 2015), taste stimulation triggered activity in a large number of regions, indicating that a large fraction of the SEZ is involved in translating gustatory information into behavior. We have therefore generated the first activity-based functional atlas of the SEZ. This atlas recapitulates anatomical and functional properties of known neuronal populations and projection fields and allows us to capture and analyze the activity of uncharacterized SEZ regions upon gustatory food stimulation in animals in different internal states. The use of this atlas should catalyze future studies aimed at investigating the function of different parts of the SEZ (Supplementary Data 1).

### Metabolic state affects taste processing in a yeast stimulus-specific manner

We then used the SEZ atlas to determine how metabolic state alters neuronal activity that ultimately will influence feeding decisions. Internal states could either influence food choice by altering how specific neurons process taste information, or they could alter chemosensory processing across a large set of circuits. Furthermore, internal states could change chemosensory processing for all stimuli or, alternatively, specifically modulate the processing of food stimuli containing particular nutrients. To assess the extent of state-dependent modulations of gustatory processing in the SEZ, we first quantified the number of active regions during food stimulation in fully fed and protein-deprived animals and compared the number of active regions across different metabolic states and food stimuli (Figure 3a). In both fully fed and protein-deprived flies, very few regions showed activity in the absence of any gustatory stimulation. While water and sucrose stimulation induced activity in multiple regions, the number of active regions did not change across metabolic states. This stands in contrast to yeast stimulation, where the number of active regions increased significantly in flies deprived of protein. Therefore, both in our imaging and in our behavioral experiments, we see clear yeast stimulus-specific changes upon protein deprivation (Figure 1a and S4).

**Figure 3.**
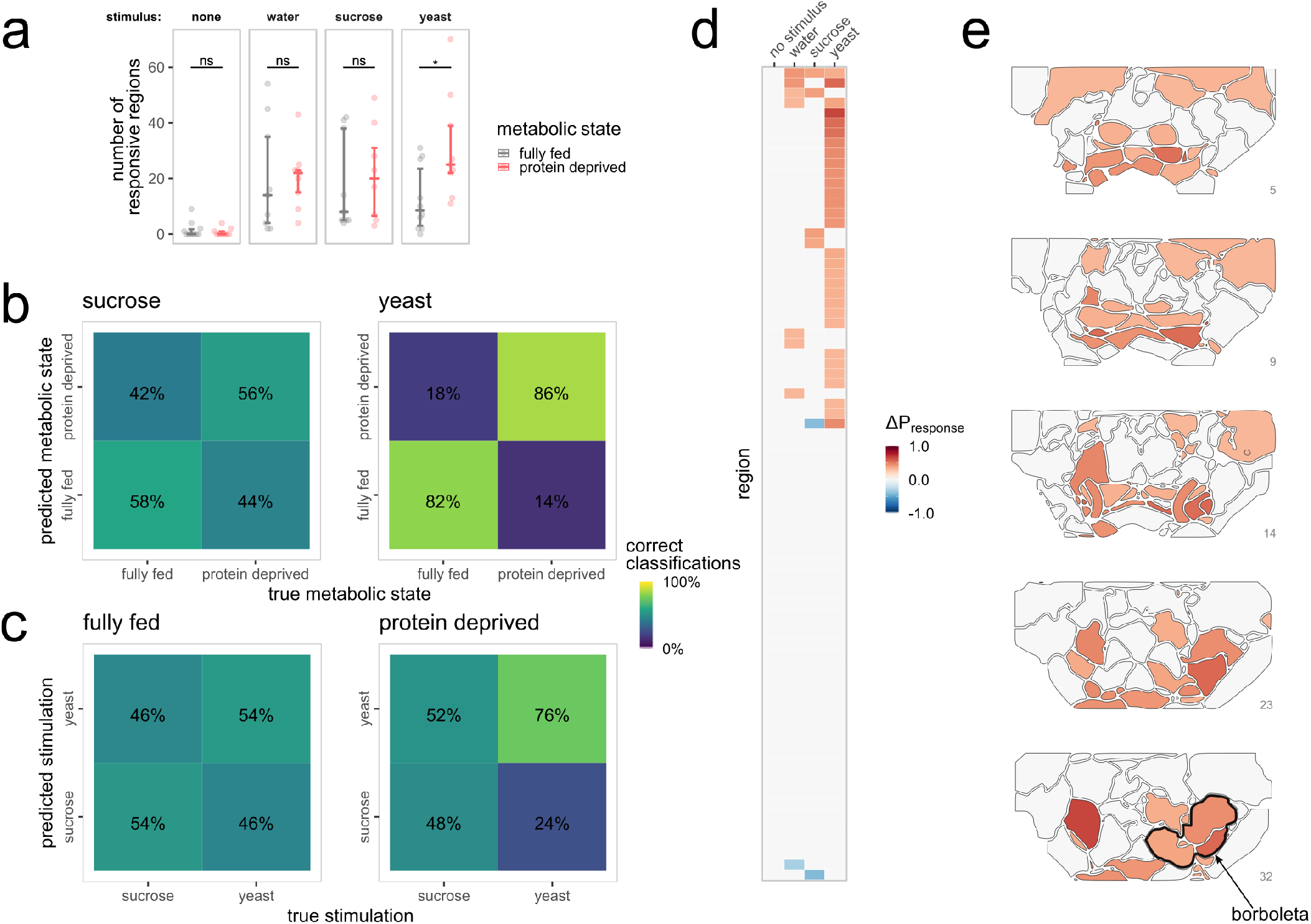
Metabolic state selectively changes taste processing in the SEZ. **a**, The number of SEZ regions responding to stimulation with different food taste stimuli. Points indicate the number of responsive regions in each experiment. Bars indicate median values, whiskers extend from first to third quartile. Wilcoxon rank-sum test, n = 8-10, ns P > 0.05, * P < 0.05. **b** and **c**, Confusion matrices showing the percentage of correct response classifications performed on activity patterns across SEZ regions. **b**, Percentage of correct metabolic state classifications using the activity pattern observed upon sucrose (left) or yeast (right) stimulation. **c**, Percentage of correct stimulus classifications using the activity pattern observed in fully fed (left) or protein-deprived (right) animals. **d**, Heatmap illustrating the change in response probability across SEZ regions upon protein deprivation for different food taste stimuli. **e**, Changes in response probability towards yeast stimulation mapped onto the SEZ atlas. The dark line highlights the borboleta outline. Only changes in response probability that exceeded ± 0.3 are plotted in **d** and **e**.

We then tested if the stimulus-induced SEZ response patterns were predictive of the current metabolic state of the animals. To this end, we trained a support vector machine (SVM) classifier on 80 % of the data and performed cross-validation using the remaining 20 %. The classifier performed at chance level when trained on the sucrose-induced response patterns (56 % and 58 %, Figure 3b). This supports our finding that changing internal amino acid state does not significantly alter sucrose taste processing (Figure 3a). When we performed the same analysis on yeast stimulus-induced activity patterns, however, the classifier performed well above chance level, predicting the correct metabolic state in 82 % and 86 % of the cases, respectively (Figure 3b). This suggests that metabolic state can be read out at the level of food-induced SEZ response patterns. Importantly, this is only possible upon stimulation with the food containing the depleted nutrient. This provides a possible neuronal correlate for the specific effects of nutrient deprivation on feeding decisions.

One way in which internal states might alter food choice is by selectively enhancing the discriminability of foods containing specific nutrients. To test this hypothesis, we trained an SVM to discriminate between different food stimuli in fully fed and protein-deprived flies. Strikingly, while the SVM performed at about chance level in fully fed animals (Figure 3c), protein deprivation increased the correct classification rate for yeast versus sucrose well above chance level. Thus, metabolic state alters the representation of relevant food stimuli in a strong and specific way.

### Metabolic state acts at different layers of taste processing

To identify which SEZ regions are modulated by internal state, we calculated how metabolic state altered the probability of each region to respond to the different food stimuli. As expected from our previous data, protein deprivation had almost no effect on the probability of regions to show a response upon water and sucrose stimulation (Figure 3d). However, nearly half of the SEZ regions in our atlas showed a clear increase in the probability of responding to yeast. Mapping these results onto the SEZ atlas revealed that protein deprivation increased the yeast response probability of regions at different levels of processing (Figure 3e). Interestingly, the few regions that showed a change in response probability upon water and sucrose stimulation showed very little overlap with the yeast stimulus regions (Figure 3d, e and S5).

Not unexpectedly, the regions showing an increased response probability upon yeast stimulation included sensory and motor regions (Figure 3e; compare Figure 2c). However, they also included unknown regions that had not been previously described and did not resemble any known taste sensory or motor neuron innervation fields. One such region showing particularly strong modulation by metabolic state is located in the deep posterior SEZ (Figure 3e). Because of its characteristic butterfly shape we refer to this region as the “borboleta”.

### Metabolic state strongly modulates the borboleta regions

To gain more insights into which regions are modulated and the nature of the modulations, we took a closer look at the responses in each region across different metabolic states. Protein deprivation resulted in a marked increase in response magnitude across SEZ regions specifically upon yeast stimulation (Figure 4a). Among the regions that showed the strongest increase in yeast responses following protein deprivation were the borboleta regions (Figure 4b-d, regions 57, 60, 9, 21, 62, 63). This increase mirrors the increase in yeast response probability we had previously observed (Figure 3e). Among all 81 atlas regions, the borboleta showed some of the strongest modulations of yeast stimulus responses (arrowheads in Figure S6a). Moreover, this modulation was stimulus-specific, as it was not observed upon sucrose stimulation (Figure S6b). We have therefore identified new SEZ regions that exhibit strong, reproducible, specific, and internal state-modulated yeast signals. Given their anatomical location, these regions are likely to receive sensory information and communicate it to the proboscis motor output, providing a possible site for internal state-dependent integration and processing of gustatory food cues informing food choice.

**Figure 4.**
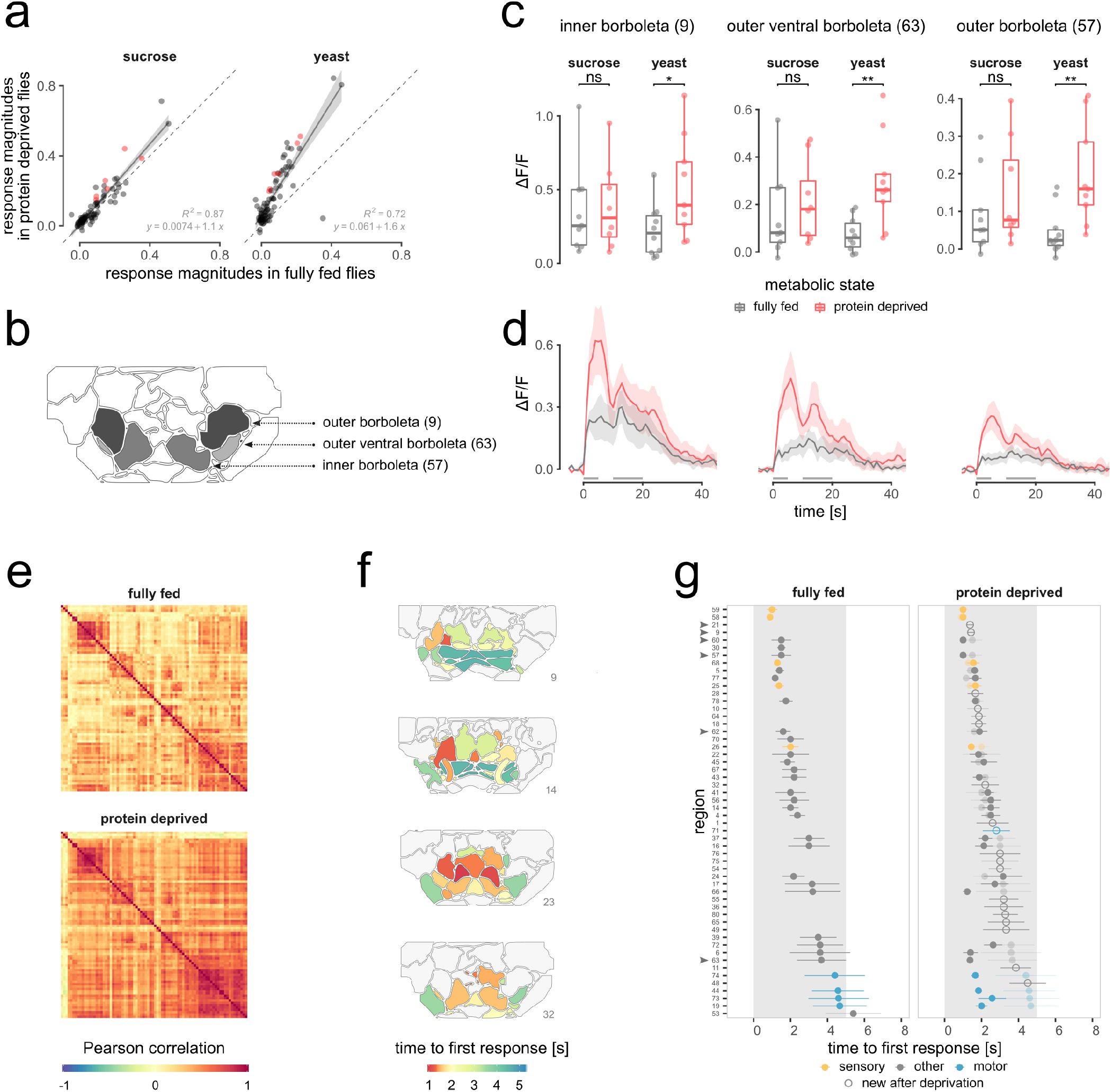
Metabolic state modulations happen at different levels of processing and increase the efficiency of motor neuron recruitment. **a**, Scatter-plots of average response magnitudes per SEZ region in fully fed vs protein deprived flies upon stimulation with sucrose (left) or yeast (right). Red points indicate the borboleta regions, dashed lines indicate the unity line, solid lines depict the regression line, and shaded areas indicate the 95 % confidence region. y indicates the fitted linear regression model, R^2^ indicates the coefficient of determination. **b**, Atlas regions constituting the borboleta (with region number in bracket). **c**, Boxplot comparing response magnitudes in the borboleta regions upon sucrose or yeast stimulation in different metabolic states. Points indicate ΔF/F response values in each experiment. Boxplots indicate median, first, and third quartile, whiskers extend to the lowest and highest values that lie within 1.5 of the inter-quartile range of the box. Wilcoxon rank-sum test, n = 9-10, ns P > 0.05, * P < 0.05, ** P < 0.01. **d**, Average traces from borboleta regions plotted in (**c**) upon yeast stimulation in fully fed (*gray*) and protein-deprived animals (*red*). Shaded area indicates mean ± sem, gray dashes indicate stimulation, n = 9-10. e, Correlation matrices depicting the Pearson correlation of yeast responses across all SEZ regions in fully fed (top) and protein-deprived animals (bottom). Matrices are clustered according to the responses in protein-deprived animals. **f**, Time to average yeast stimulus-induced response onset in fully fed flies visualized across the functional SEZ atlas. **g**, Average time to response onset upon yeast taste stimulation in fully fed (left) and protein-deprived animals (right). Only regions with responses in ≥ 5 animals are shown. Points and line range indicate mean ± sd, n = 5-9. Colors indicate region categories (sensory yellow, motor blue), the gray shaded area indicates the stimulation time, open points indicate regions only active in protein-deprived flies, transparent points indicate data from fully fed flies plotted in the protein-deprived graph for ease of comparison.

### Response dynamics change with metabolic state

How might the increase in response probability and strength induced by metabolic state lead to changes in food preference? We hypothesized that these changes could increase the efficiency of signal propagation both at the temporal and circuit level, leading to a stronger and faster recruitment of SEZ regions. We therefore examined the correlations between yeast stimulus-induced response time series across atlas regions, and how these correlations were altered by internal state. We found that prominent clusters of highly correlated regions present in fully fed flies were maintained after protein deprivation (Figure 4e). However, protein deprivation resulted in increased correlation across large parts of the network, including new regions which were previously weakly correlated. This supports the hypothesis that nutrient deprivation enhances signal propagation within a core network while simultaneously recruiting new regions. This highlights the broad contribution of the SEZ network to internal state-dependent sensorimotor processing and suggests that metabolic state not only increases the strength of the sensory input but also acts on multiple regions of the SEZ to increase protein appetite.

Next, we focused on the temporal emergence of food-induced activity across the SEZ. For this, we calculated the time at which we observed the first response upon sensory stimulation in each region of the SEZ (Figure 4f). As expected, sensory regions were the first to respond upon stimulation, whereas motor regions were the last, with the remaining regions responding in between (Figure 4f and g). Protein deprivation had two effects. As expected from the above results, upon protein deprivation new regions showed yeast-evoked activity. Furthermore, there was a striking consistent decrease in response times in motor regions upon protein deprivation (Figure 4g). This stands in contrast to sensory regions, where metabolic state did not alter response timing. Thus, metabolic state results in drastically reduced response time of the motor neuron regions, which is likely to significantly increase the probability that the animal will initiate and sustain feeding on a specific food source, thus biasing its food choice.

### Mating and reproductive states are synergistically integrated at the motor neuron level

Mating and protein deprivation synergistically increase yeast feeding in females (Figure 1a). By doing so the fly anticipatorily increases protein intake to prevent a lack of building blocks required for oogenesis. To visualize where in the SEZ reproductive state modulates yeast responses, we calculated the change in yeast responses upon mating in yeast-deprived females. While the changes observed upon mating were rarely statistically significant (Figure S7), the regions that showed the biggest changes in responses to yeast stimuli mapped to putative motor output regions (Figure 5a and S7). This stands in contrast to metabolic state which leads to strong and statistically significant changes in yeast responses throughout the SEZ (Figure S6).

**Figure 5.**
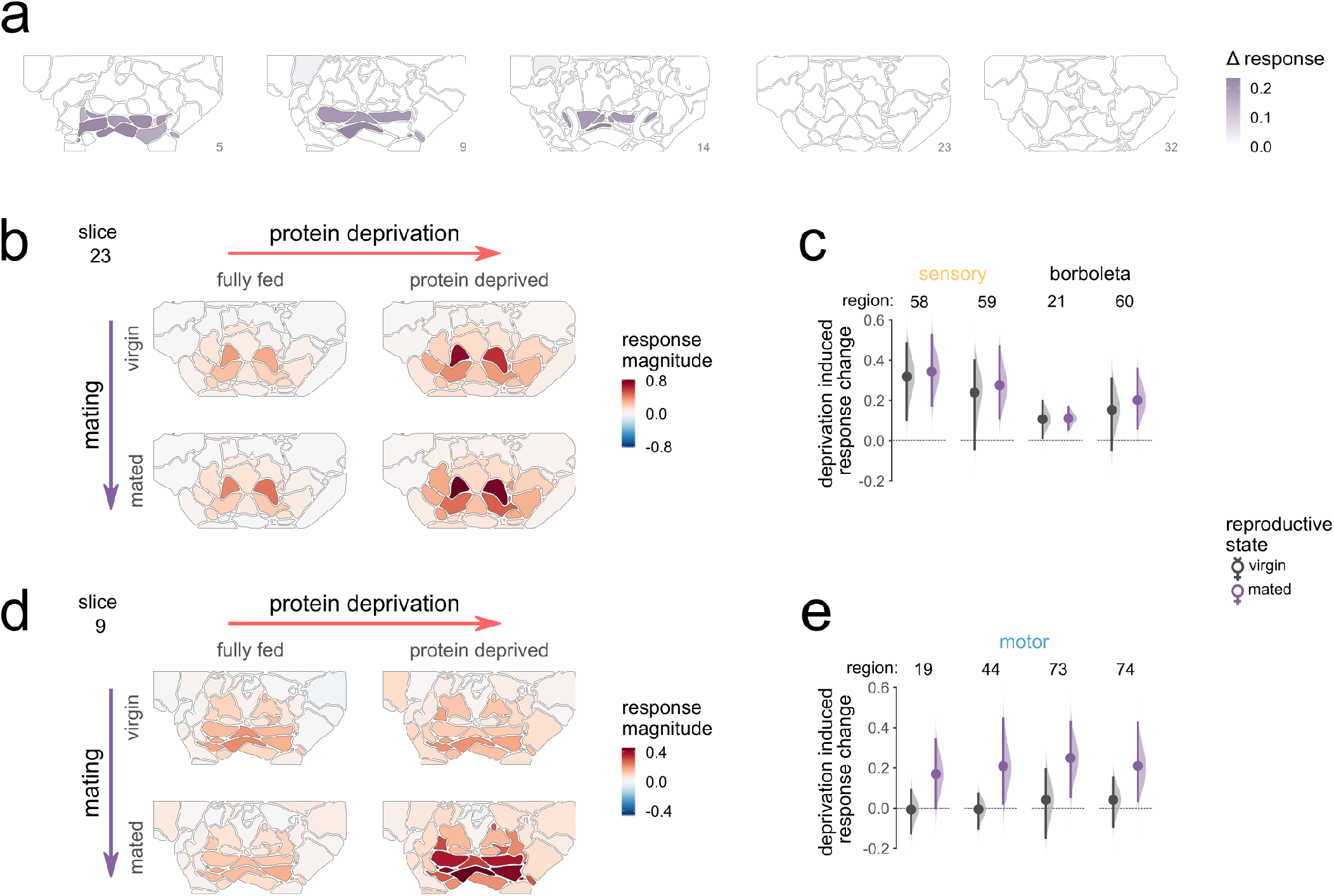
Reproductive state acts on motor neuron regions. **a**, Magnitude of the changes in average response upon mating in different SEZ atlas regions. The shaded regions are the ten regions showing the strongest or significant changes of all regions plotted in Figure S7. **b**, Atlas slice 23 containing sensory regions, with yeast stimulus elicited response magnitudes plotted for the four different internal states. **c**, Bootstrapped mean differences in response magnitude upon protein deprivation, for virgins, *black* and mated females, *purple*. Plotted for regions 58 and 59 (sensory) and 21 and 60 (borboleta). Points indicate the mean of bootstrapped differences of means between fully fed and deprived flies. Bars indicate confidence intervals of the bootstrapped distributions. Shaded regions indicated density plots of the bootstrapped distributions. **d**, like **b** but for atlas slice 9 containing motor regions. e like c but for motor regions (19, 44, 73, 74). See Figure S8 for the remaining atlas slices.

Given that the two states act synergistically on food choice, we reasoned that the effect of mating should be best visible when analyzing both states together. We therefore compared the magnitude of the change in yeast-stimulus induced activity upon protein deprivation in virgin and mated females. In sensory and borboleta regions, protein deprivation led to strong changes in response, independently of the mating state (Figure 5b, c and S8). Protein deprivation, however, had no noticeable effect on yeast-induced motor neuron region activity in virgin females (Figure 5d). Metabolic state mainly affected yeast-induced activity in the motor neuron region in mated females (Figure 5d and e). This shows that reproductive and metabolic state synergistically act on the feeding motor neuron regions to anticipatorily alter food choice and enhance the reproductive success of the female. Different internal states therefore synergistically shape feeding decisions by acting at different levels of the sensorimotor loop using a mix of global and local modulatory strategies.

### The outer borboleta region is sufficient to induce protein appetite

One of the major challenges in studying neuronal processing across the brain is the difficulty in functionally validating the behavioral relevance of observed modulations. The neurogenetic tools available in *Drosophila*, coupled with comprehensive databases that map these tools to specific brain regions have revolutionized our ability to interrogate circuit function in this system. Having identified the borboleta region as a node of convergence between sensory input and metabolic state, we set out to develop a strategy to test the behavioral relevance of this region in mediating yeast preference. This required us to efficiently query the databases of genetic driver lines for expression in the SEZ regions in our functional map. To do so, we performed a bridge registration between our physiological *in-vivo* SEZ standard and the commonly used JFRC2 anatomical standard brain (Jenett et al., 2012) (Figure 6a, see also Materials and Methods for details) allowing us to intersect our data with data from the Braincode database (Panser et al., 2016). Braincode performs spatial clustering of the *Drosophila* brain based on the expression patterns of thousands of Gal4 driver lines. We quantified the overlap between borboleta regions and Braincode clusters and identified Braincode clusters 11 and 36 to have the strongest overlap with outer and inner borboleta, respectively (Figure 6b). Using the Braincode database, we then selected sparse Gal4 lines which showed expression in these clusters for functional validation (Figure 6b, c and S9). Intersecting available 3D expression data confirmed that these lines had overlapping expression only in the outer or inner borboleta regions and nowhere else in the brain (Figure 6c).

**Figure 6.**
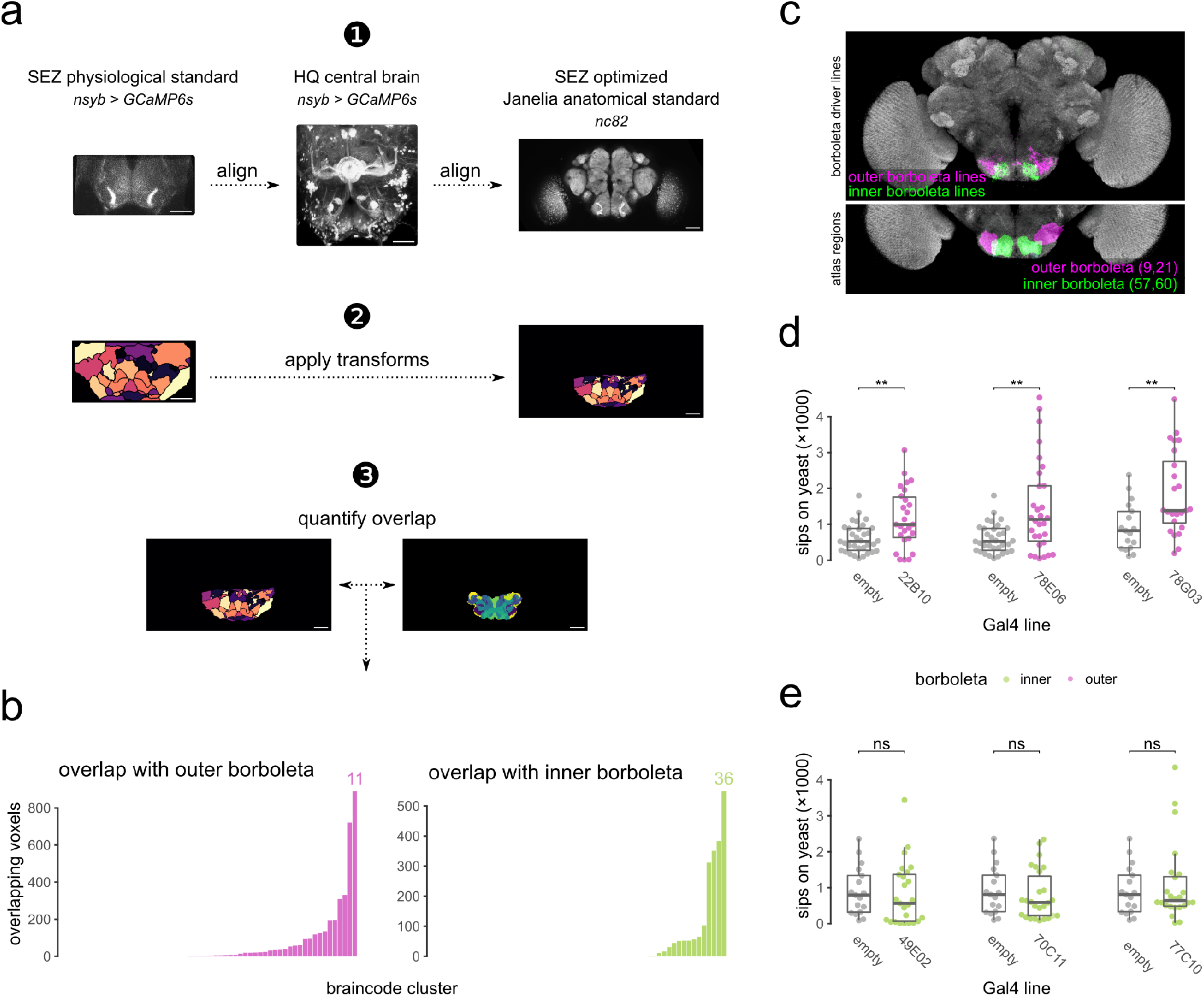
Outer borboleta neurons drive protein appetite. **a**, Schematic illustrating the bridge registration steps and the intersection with the Braincode atlas. (1) Our SEZ standard was aligned to the fixated JFRC2 standard brain *via* a bridge registration to a high-resolution scan of the central brain. (2) Transforms from (1) were applied on our SEZ atlas to transfer it from *in-vivo* SEZ space to fixated JFRC2 space. (3) We quantified the overlap of borboleta regions and Braincode clusters to select candidate Gal4-driver lines from the Janelia FlyLight collection. **b**, Quantification of the overlap between SEZ and brain code atlases identifies Braincode cluster 11 as showing the largest overlap with the outer borboleta (left, *magenta*) and brain code cluster 36 as showing the largest overlap with the inner borboleta (right, *green*) **c**, Overlap of the selected outer borboleta (*magenta*) and inner borboleta (*green*) driver-lines within the Janelia anatomical standard brain JFRC2 (z-section 75-150) and the corresponding SEZ atlas regions (outer borboleta in *magenta* and inner borboleta in *green*) below for comparison. **d** and **e**, Number of sips as measured by the flyPAD assay from 10 % yeast by mated female fully fed flies maintained for 2h at restrictive temperature. The genotype of the Gal4 lines used to drive TRPA1 expression is depicted below the plots with empty depicting the matching empty-Gal4 control. Results for lines showing expression in the outer borboleta are plotted in **d** and for lines showing expression in the inner borboleta are plotted in e. Points indicate the average number of sips on yeast for each fly. Boxplots indicate median, first, and third quartile, whiskers extend to the lowest and highest values that lie within 1.5 of the inter-quartile range of the box. **d** Wilcoxon rank-sum test, n = 17-33, ** P < 0.01. **e** Wilcoxon rank-sum test, n = 17-26, ns P > 0.05.

To probe a possible causal relationship between neurons innervating the borboleta regions and protein appetite, we used thermogenetic activation of the labeled neurons in combination with the flyPAD feeding assay (Itskov et al., 2014). Strikingly, in fully fed flies, conditional activation of driver lines targeting the outer borboleta using our protocol increased feeding on the yeast food spot when compared to control flies (Figure 6c and d). We validated these results using a separate tracking-based assay (Corrales-Carvajal et al., 2016) (Figure S10). Conversely, experiments using driver lines with expression in the inner borboleta region revealed no yeast feeding phenotypes (Figure 6c and e). We note that in some replicates, activation of neurons labeled in the line 22B10 led to a reduction in yeast feeding, which is consistent with these neurons having a function in mediating yeast feeding and suggests that artificial activation of these neurons may also lead to loss-of-function phenotypes. Importantly, experiments with lines labeling neurons innervating the outer borboleta did not reveal increases in sucrose feeding (Figure S11). This parallels the absence of internal state-modulated sucrose responses in this region and suggests that neurons therein specifically alter yeast appetite.

Flies adapt their food intake by modulating the feeding microstructure (Itskov et al., 2014). Protein deprivation increases the probability to initiate a feeding burst on yeast (quantified as a decrease in the inter burst interval) and the duration of the feeding bursts (quantified by the number of sips per burst) (Figure S12a and b). These two parameters are thought to be regulated independently: The initiation of feeding bursts is thought to be mostly modulated by inputs from bristle gustatory neurons, while the feeding burst length seems to depend on taste input from peg gustatory neurons (Steck et al., 2018). To test if the outer borboleta regulates yeast feeding by modulating sensory input from specific taste neurons, we inspected the effect of activating outer borboleta neurons on the feeding microstructure. In two of three lines, activation of neurons that labeled the outer borboleta led to a strong decrease in the intervals between yeast feeding bursts (Figure S12c), without affecting the length of these feeding bursts (Figure S12d). In the third line, the control had a short inter-burst interval, likely precluding an observable effect. These data suggest that the outer borboleta has a specific role in increasing the probability to initiate feeding on yeast, rather than in sustaining it. Furthermore, these data lend further support to the idea that different aspects of the feeding microstructure are processed by separate circuits in the SEZ, and suggest that the outer borboleta specifically modulates sensory information coming from the bristle sensillae. Given that artificial activation of these neurons does not fully recapitulate the changes in feeding microstructure observed upon protein deprivation, these results also suggest that we are not simply activating neurons that broadcast the metabolic state to the rest of the SEZ. Taken together, these data show that by bridging our pan-neuronal imaging data to lines labeling specific modulated regions in the atlas, we are able to identify a region that plays a causal and selective role in driving food choice.

## Discussion

Here, we used pan-neuronal imaging of the ventral part of the female *Drosophila* brain to show that metabolic state modulates taste processing at global scales to shape food choice. Our observation that gustatory stimulation leads to the recruitment of a large fraction of SEZ neurons echoes recent findings using different sensory stimuli in different species. A study pioneering the use of pan-neuronal imaging in flies observed that sweet and bitter stimuli recruit a large set of mostly separable neuronal populations in the SEZ (Harris et al., 2015). While the focus on imaging somata did not allow the authors to dive deeper into the mechanistic relevance of this observation, this study and more recent studies using auditory stimuli in flies (Pacheco et al., 2021), as well as visual and olfactory stimuli in fishes and rodents (Allen et al., 2019; Portugues et al., 2014), confirm that sensory representations encompass larger fractions of brains than classically assumed. This is not surprising given that sensory information is used to inform many different computations. While classically such computations were often thought to be implemented in a linear and sequential fashion, recent studies suggest that computations underlying decision-making and behavior are performed in a distributed and highly parallelized manner (Suri et al., 2020). This likely leads to a broad recruitment of neuronal populations upon sensory stimulation. The ability to image neuronal dynamics broadly across the brain allows for new perspectives on how sensory information is used by neuronal systems to generate decisions such as choosing what food to eat.

We find that the metabolic state of the animal strongly modulates taste processing across a large portion of the *Drosophila* brain. Importantly, these modulations are food-type specific. As such, we find that protein deprivation specifically increases neuronal activity in large parts of the SEZ only when the animal experiences protein-rich food and not sucrose. Our findings are in agreement with the few classic studies in primates documenting that in the orbitofrontal cortex (OFC) neuronal responses to the sight, smell, or taste of food are modulated in a satiety-specific way (Rolls, 2015). These reductions in sensory-induced neuronal activity are extremely specific, as only responses to sensory stimuli associated with the ingested food were found to be quenched. This modulation of taste-induced activity in the OFC has also been observed in humans (Kringelbach et al., 2003) and is thought to link the valuation of specific foods to their sensory representations. These changes in neuronal representations are thought to mechanistically underpin the phenomena of “alliesthesia” (Cabanac, 1971) and “sensory-specific satiety” (Rolls et al., 1981), phenomena that are thought to play important roles in shaping food choice in humans. However, this mechanistic hypothesis remains to be tested. Recently, it has also been reported that metabolic states such as thirst and hunger alter putative representations of arbitrary sensory stimuli paired with food or water rewards in different brain regions (Allen et al., 2019; Burgess et al., 2016; Livneh et al., 2017, 2020). Given that these changes in representations correlate with the pleasantness rating given to the foods, the willingness of the satiated animal to consume these foods, or their performance in decision-making tasks, they are thought to be likely functionally relevant. Our finding that activating neurons in the borboleta region biases food choice towards protein-rich food, provides first evidence that the changes in representation upon alterations in internal state functionally underlie decision-making.

The SEZ plays a key role in mediating sensorimotor transformations in insects. This is best appreciated in the case of taste and feeding, as all mouthpart gustatory and motor neurons project to and from the SEZ. While the molecular and anatomical logic of gustatory processing at the primary sensory level is relatively well understood (Scott, 2018), we surprisingly lack an understanding of the circuitry and computational principles by which taste information is processed throughout the brain. This stands in stark contrast to other senses such as olfaction where we have a solid grasp of these processes up to brain centers that mediate learning and memory (Schlegel et al., 2020). Moreover, akin to the vertebrate brainstem, the insect SEZ is emerging as the neuropil managing a large part of motor transformations shaping motor outputs in insects (Gal and Libersat, 2006; Knebel et al., 2019; Pulver et al., 2015). Nevertheless, our understanding of the SEZ remains rudimentary. A key challenge is the lack of clear anatomical structural subdivisions in the SEZ. As we show, our functional SEZ atlas provides such a structural organization. It encompasses most known anatomically identified projection groups and expands these by new regions which have not been previously explored. Importantly, we show that the atlas allows for the formulation of functional hypotheses, which can be tested experimentally by bridging the atlas to existing transgenic expression lines. Similar to previous findings in vertebrates (Allen et al., 2017) we find that not all sensory-modulated regions lead to behavioral phenotypes when perturbed. This suggests that while neuronal representations can be widely distributed, behavior is likely to only be influenced by perturbations of specific regions. Whether these regions also constitute important functional processing units under physiological conditions will be an important point for future studies aimed at bridging brainwide activity mapping with functional circuit dissections.

The functional similarities between the SEZ and the vertebrate brainstem are striking, particularly with respect to the regulation of feeding. Both regions harbor the primary gustatory centers (Vendrell-Llopis and Yaksi, 2015; Yarmolinsky et al., 2009), centers generating the feeding motor program (McKellar, 2016; Moore et al., 2014), as well as nuclei modulating food intake according to internal states (Carter et al., 2013; Jourjine et al., 2016; Marella et al., 2012; Nectow et al., 2017; Yapici et al., 2016). Our work suggests that these three distinct units are functionally connected within the SEZ, with the metabolic state either acting directly on them or even being generated within the circuit. We also identify the “borboleta” regions, and more specifically neurons with arborizations within a particular subregion, as key in orchestrating protein appetite. We clearly show that this region is modulated by both sensory stimuli and metabolic states. How neurons within the outer borboleta mediate protein appetite will be an important avenue for future research. Our data are consistent with a model in which sensorimotor transformations within the SEZ account for most of the observed adaptation of feeding to the nutritional needs of the animal. Current models of the vertebrate brainstem, however, treat the gustatory information stream modulating feeding as separate from those that mediate the internal state and the feeding motor program. As discussed above, internal states are thought to alter gustatory representations at the level of the insula or the OFC, which then relay the food intake decision to the brainstem. Such a model is also supported for protein homeostasis in humans, where protein-deficient subjects were found to have increased activity in the OFC upon exposure to sensory properties of protein-rich foods, but not other foods (Griffioen-Roose et al., 2014). Based on our data, we propose that gustatory information reaching the NST in the brainstem might also be modulated by brainstem circuits signaling the internal state of the animal and impinge more directly on premasticulatory circuits to alter feeding. Recent findings linking circuits that convey internal state information to IRt/PCRt (Nakamura and Nakamura, 2018), as well as classic studies showing changes in the representation of gustatory information at the level of the NST in rats (Giza and Scott, 1983) support such a model. Local computations in the brainstem or the SEZ could act in parallel with distributed computations including the OFC, or higher-order fly brain regions, to coordinate feeding with higher-level foraging decisions, such as the decision to continue eating from a food patch or plate or to leave it in search of other, more nutritionally fitting food locations.

We find that metabolic state alters taste representations in a sensory-specific but anatomically distributed manner across large portions of the fly ventral brain. These results echo the finding that in mice, thirst globally alters sensory representations (Allen et al., 2019). It remains unclear, however, if other internal states also act on behavior by altering sensory representations over large regions or whether they act in more focused ways. This question is particularly relevant when considering how animals integrate multiple ongoing states. We have a very poor understanding of how such state integrations are implemented, despite their clear ethological relevance, given that animals in natural habitats must consider multiple ongoing internal states when deciding which behavioral decisions to make. We show that in the female fly, reproductive state has a fundamentally different mode of action on gustatory representations than metabolic state. Supporting earlier findings made in specific gustatory neurons (Steck et al., 2018), mating barely altered taste representations across most of the SEZ. The effect of mating was extremely specific for putative motor neuron regions. Mating increased activity observed in regions corresponding to motor neuron innervation fields upon yeast taste stimulation. Importantly, this was only evident in protein-deprived females, providing a neuronal explanation for the synergistic effect of both states on protein appetite. Our data suggest that mating acts in a focused way to increase the likelihood that relevant food stimuli will trigger a feeding motor output. This localized effect of reproductive state on gustatory representations contrasts strongly with the broader effect observed upon alterations in metabolic state. Why might metabolic and reproductive state modulate food choice in such different ways? First, changes in metabolic state are gradual and are known to affect many other processes such as learning and memory (Krashes et al., 2009). As such a broader and more distributed neuronal implementation of valuation by metabolic state would allow for the modulation of a variety of computations. Second, nutrient levels can drop to threatening levels in virgins, which requires that the animal is able to detect metabolic demand independent of reproductive state. By strategically increasing the likelihood that gustatory information triggers a feeding motor output, reproductive state synergistically amplifies the gustatory and metabolic state information without altering the representation of metabolic state itself.

The regulatory logic we have identified also implements a predictive change in food choice that anticipates the future nutritional needs of the female. By amplifying states of low metabolic deficit, which in themselves do not pose a challenge to the virgin female, mating increases food intake of reproductively relevant nutrients before a functionally relevant nutritional deficit manifests. At the circuit level, we have previously shown that anticipatory appetite for salt and protein are induced by the canonical SPSN/SAG mating circuit, which relays information about mating state from the reproductive tract to the brain and shapes how gustatory information is processed by the animal (Walker et al., 2015). While the circuitry linking SAG neurons to the feeding motor output remains to be elucidated, our data suggest that circuits downstream of SAG neurons specifically gate how sensory information is translated into a motor output. In the case of receptivity, SAG downstream neurons act on descending premotor interneurons to gate whether sensory information is translated into the corresponding motor neuron output (Wang et al., 2021). Our findings therefore suggest a coherent picture in which the post-mating circuitry alters behavior in mated females by controlling the transformation of sensory information late at the motor output level. The specific case of food choice is special in this context, as it requires the additional integration of metabolic state information to allow for a predictive alteration in how taste information is processed.

Our data suggest a coherent picture of how internal states shape nutrient-specific appetites and thus food choices. Deficiencies of specific nutrients lead to increased responsiveness of taste circuits detecting foods containing those nutrients (Steck et al., 2018). The additional recruitment of downstream circuits, such as neurons innervating the outer borboleta, leads to an increased discriminability of the sensory properties of the required foods and a more efficient activation of downstream circuits. This ultimately increases the probability and the efficacy by which the feeding motor output is initiated and maintained. This is reflected in a clear reduction in the time required for a sensory stimulus to initiate feeding motor neuron activity. This increase in the probability and efficacy of initiating feeding from a given food source is specific in two ways. First, we observed little change in the probability of sucrose or water stimuli to recruit SEZ regions upon protein deprivation. This strengthens the concept that the representation of different chemosensory modalities is separate at the circuit level. As such, activating neurons in the outer borboleta only increases yeast feeding, but not sucrose feeding. Second, our data suggest that different aspects of the feeding microstructure are regulated by separate sensory circuits. While our previous data showed that taste pegs specifically control the maintenance of a feeding burst, here we show that the activation of neurons in the outer borboleta specifically increases the likelihood that the animal will initiate yeast feeding. The separation of resource specificity and motor program modulation allows the parallel but separate implementation of valuation computations within the SEZ. This enables a differentiated and graded adjustment of food choice to meet the current and future needs of the animal. We therefore identify a generalizable principle by which internal states can be integrated to shape sensory-guided decisions and propose a strategy by which similar principles can be efficiently uncovered and functionally validated for other internal states and decisions.

## Supporting information

Supplemental Data 1 - SEZ Atlas

Supplemental Data 2 - SEZ Template

Supplemental Video 1 - Example Recording

## ACKNOWLEDGMENTS

We thank Célia Baltazar and Lúcia Serra for technical assistance. We thank Alexandre Laborde, Aaron Ostrovsky, João Baúto and the Champalimaud Software Platform for assistance in establishing the registration workflow on the HPC cluster. We thank the Champalimaud Fly Platform for support with acquiring anatomical images. We thank Terufumi Fujiwara for sharing fly lines. We thank Gil Costa for assistance with illustrations. We thank Ricardo Henriques for creating the L^A^T_E_X template. We thank Ibrahim Tastekin, Samuel Walker, Michael Orger, Eugenia Chiappe and the whole Behavior and Metabolism laboratory for many fruitful discussions, valuable feedback throughout the project and comments on the manuscript. Flies obtained from the Bloomington Drosophila Stock Center (NIH P40OD018537) were used in this study. D.M. was supported by a DFG Research Fellowship (MU 4116/1-1). D.G. was supported by a doctoral fellowship PD/BD/114273/2016 from the Portuguese Foundation for Science and Technology (FCT). The project leading to these results has received funding from “la Caixa” Banking Foundation to C.R. under the project code LCF/PR/HR17/52150002, a grant funded under the Lisbon Regional Operational Programme (POR) by national funds through the FCT - Fundação para a Ciência e Tecnologia, I.P. and co-financed by the European Regional Development Fund (ERDF) under the project 02/SAICT/300081/2017, and supported by the research infrastructure Congento, co-financed by Lisboa Regional Operational Programme (Lisboa2020), under the PORTUGAL 2020 Partnership Agreement, through the European Regional Development Fund (ERDF) and FCT - Fundação para a Ciência e Tecnologia (Portugal) under the project LISBOA-01-0145-FEDER-022170. Research at the Centre for the Unknown is supported by the Fundação Champalimaud.

## Materials and Methods

### Fly husbandry & internal state manipulations

Flies were reared on yeast-based medium (per liter of water: 8 g agar [NZYTech, Portugal], 80 g barley malt syrup [Próvida, Portugal], 22 g sugar beet syrup [Grafschafter, Germany], 80 g cornflour [Próvida, Portugal], 10 g soya flour [A. Centazi, Portugal], 18 g instant yeast [Saf-instant, Lesaffre, France], 8 mL propionic acid [Argos], and 12 mL nipagin [Tegospet, Dutscher, UK] [15 % in 96 % ethanol] supplemented with instant yeast granules on the surface [Saf-instant, Lesaffre, France]) and kept at 25° C, 70 % relative humidity in a 12 h / 12 h light-dark cycle. Flies were reared on YBM and transferred to fresh YBM on days 0 and 2 after hatching. Deprivation was induced by transferring 3 days old flies to vials containing a half paper tissue soaked with 5 mL of 100 mM sucrose solution for 10 days. Fully fed flies were age-matched. All flies were transferred to vials with the respective fresh medium every two days. For the mated conditions 10 female flies were kept together with 5 male flies. For virgin conditions, 10 virgin flies without males were kept per vial.

### Fly stocks & genetics

All calcium imaging experiments were performed with female flies of the geno-type: *w*; 20×UAS-GCaMP6s / +; 57C10-Gal4 / +*, expressing UAS-GCaMP6s under the control of the pan-neuronal *nSyb-Gal4* driver line *57C10-Gal4*. Flies of this genotype were used to perform the flyPAD food choice experiments shown in Figure 1a and S12a, b. Thermogenetic activation experiments in Figure 6 and Supplementary Figures S10, S11 and S12c, d were performed with flies of the genotype *w^1118^ ; UAS-TrpA1 / +; X-Gal4 / +*, expressing the temperature-sensitive cation channel dTRPA1 under the control of different specific Gal4 driver lines from the Janelia FlyLight collection (where X-Gal4 is *22B10-Gal4, 49E02-Gal4, 70C11-Gal4, 77C10-Gal4, 78E06-Gal4, 78G03-Gal4* or empty-Gal4 as control). For immunostainings, the specific Gal4 driver lines were crossed with flies carrying UAS-CD8::GFP on the second chromosome.

### Fly preparation for calcium imaging

13-14 days old flies were briefly cold anesthetized on ice and glued to a custom-made acrylic block with their thorax using fast UV curing glue (Bondic, Niagara Falls, New York, US). To facilitate gustatory stimulation and to reduce brain movement, the proboscis was extended using a blunt needle (B30-50; SAI Infusion, Faridabad, Haryana, India) attached to a vacuum pump (N86KN.18; KNF DAC GmbH, Hamburg, Germany) and fixed in an extended position by carefully applying UV curing glue only to the proximal part of the proboscis using an insect pin, such that the labellum could move freely. The front legs were removed to prevent flies from touching the food stimulus during the experiment. The anterior part of the head capsule was placed through a hole in a plastic weigh boat that was fixed on top of the fly. The space between the head and the weigh boat was sealed with UV curing glue. The head capsule was covered with carbogenated (95 % O_2_, 5 % CO_2_) adult hemolymph-like saline (AHL; 103 mM NaCl, 3 mM KCl, 5 mM TES, 10 mM trehalose dihydrate, 10 mM glucose, 2 mM sucrose, 26 mM NaHCO_3_, 2 mM CaCl_2_ dihydrate, 4 mM MgCl 2 hexahydrate, 1 mM NaH_2_PO_4_, pH 7.3) and a window was cut between the eyes and the ocelli, thereby removing the antennae. Trachea covering the brain were removed and the esophagus was transected to allow for unoccluded visual access to the SEZ.

### Volumetric calcium imaging

All imaging was performed using a resonant-scanning two-photon microscope (Scientifica, UK) allowing for frame rates of ~ 60 Hz. The system was equipped with a 20x NA 1.0 water immersion objective (Olympus, Japan), controlled by a piezo-electric z-focus, allowing for fast volumetric scans. A Chameleon Ultra II Ti:Sapphire laser (Coherent, Santa Clara, CA, USA) was used to excite GCaMP6s (Chen et al., 2013) at 920 nm. Imaging data were acquired using SciScan (Scientifica, UK). 60 s recordings of the SEZ volume were performed at 1 Hz volume rate covering 512 × 256 × 60 voxels at voxel dimensions of ~0.5 × 0.5 × 3.6 µm. Scanning was performed in sawtooth mode and 5 z-planes acquired during flyback were removed. During imaging, the brain was constantly perfused with AHL saline bubbled with carbogen (95 % O_2_, 5 % CO_2_). Imaging was performed with flies at four different internal states: fully fed & virgin (n = 8 animals), fully fed & mated (n = 10 animals), protein deprived & virgin (n = 8 animals), protein deprived & mated (n = 9 animals).

### Gustatory stimulations

Glass capillaries for stimulations (GC15F-10, Harvard Apparatus, Edenbridge, Kent, UK) were pulled using a laser pipette puller (P2000; Sutter, Novato, CA, USA) to have a blunt end and an inner diameter fitting the fly proboscis. 200 mL pipette tips were cut to fit the glass capillaries and sealed with Parafilm (Amcor, Zürich, Switzerland). Capillaries were filled with taste solutions and positioned in front of the fly proboscis using a micromanipulator (Sensapex, Finland). Positioning and stimulation were performed under visual control using a PointGrey Flea3 camera and a custom Bonsai script (Lopes et al., 2015). All stimuli were prepared in MilliQ water (Merck KgaA, Darmstadt, Germany). Stimuli consisted of 200 mM sucrose, 10 % w/v yeast (Saf-instant, Lesaffre, France) and water. During imaging, two manual taste stimulations were performed at 10-15 s and 20-30 s.

### Preprocessing pipeline

Data preprocessing was performed using available command line tools and Python packages as well as custom written Python code. Several parts of the pipeline build on functions from the `nibabel` and `nilearn` Python suits. Movement correction and registration steps were performed on a computational cluster provided by the Champalimaud Software Platform.

First, we performed rigid movement correction of each 3D time series using `3dvolreg` from the `afni` toolkit (Cox, 1996). Next, to correct for shifts between recordings in the same animal, we aligned the first volume of each recording to the first volume recorded in an animal performing a non-linear registration using the antsRegistration function from the Advanced normalization Toolkit (`ANTs;` Avants et al., 2011). To have all recordings in the same 3D space, all data were aligned to a common SEZ standard (Supplementary Data 2). The SEZ template was generated with the `ANTs` script `antsMultivariateTemplateConstruction2.sh`, using 7 representative SEZ volumes that were pre-aligned using a manual landmark-based approach. Manual landmarks were set in ITK-SNAP (http://www.itksnap.org; Yushkevich et al., 2006). These landmarks were: the dorsal and ventral part of the AMS1 region and the entrance points of the pharyngeal-, the accessory pharyngeal- and the labial nerves. Pre-alignment was performed with the `antsLandmarkBasedTransformInitializer` function. Next, all data were registered to the SEZ standard by applying the same pre-alignment followed by a rigid, an affine, and finally, a non-linear (SyN, symmetric normalization; Avants et al., 2011) registration step within the antsRegistration function. Finally, we applied all concatenated transforms to all individual volumes recorded using the `antsApplyTransforms` function. Registrations were visually inspected for successful alignment as assessed by the overall overlap of the neuropil signal, major distortions and overlap of the manually defined landmarks. Data in which the quality criteria described above such as the overlap of the landmarks or the neuropil borders was not achieved were excluded from further analysis.

### Functional segmentation & signal extraction

To build the functional atlas we applied a gaussian filter and ΔF/F normalization to each individual 4D image stack (x × y × z × t). We then averaged data for each stimulus recorded in each internal state, across repetitions (flies). We used the `nilearn/scikit-learn` (Abraham et al., 2014; https://nilearn.github.io) implementation of group model Canonical ICA (`nilearn.decomposition.CanICA`; Varoquaux et al., 2010 in Python to extract 80 spatially independent components from the recordings. Within the ICA a 2 × downsampling step was performed and a manually drawn mask was applied to restrict the analysis to the brain neuropil. We ran the nilearn.regions. `RegionExtractor` function to split bilateral components into individual regions. We converged all spatial components into a binary map by assigning each voxel to the component with the highest z-score and performed several erosion and dilation steps. Finally, we used the `NiftiLabelsMasker` function to extract signals for each atlas region from each individual animal by averaging across all voxels belonging to a region. To perform the same analysis on temporally shuffled data (Figure S2c and d) we shuffled each voxel of the input data along the time dimension.

### Atlas visualization

To visualize data on the functional atlas we created a vector version of five slices (5, 9, 14, 23, 32) of the atlas that covered all 81 regions. We used a custom function in R that uses `ggplot` (Wickham, 2009) to map color-coded responses on this atlas representation.

### ΔF/F calculation and temporal alignment

Analysis of the extracted time series was performed in R (R Core Team, 2020). Data were normalized by calculating ΔF/F = (Fx-F0)/F0 with F_x_ being the fluorescence at time point x and F_0_ being the average fluorescence during 6 s directly preceding the first stimulation. To correct for variations in the manual stimulation between animals, in each animal, the response onset in the left PMS4 sensory region was defined as time-point 0. Response onset was defined as the time point where fluorescence exceeded 4 × the SD of the background activity, with background SD quantified during 8 s before first stimulation.

### Responsive regions and response probabilities

We defined a responsive region as a region in which the response during the first 5 s stimulation exceeded a threshold of 4 × the SD of the background activity, with background activity calculated from 6 s preceding the first stimulation. To calculate the response probability for each region in each internal state we divided the number of detected responses by the number of total recordings. To calculate the change in response probability upon protein deprivation, we subtracted the response probabilities in fully fed flies from those in protein-deprived flies.

### Response pattern classification

Pattern classification in Figure 3b and c were performed on response magnitudes (see Response quantification section). We trained a support vector machine (SVM) classifier using the svm function of the e1071 package in R. We used ~ 80 % of the flies as training dataset and tested on the remaining ~ 20 % (2 out of 8-10 flies for Figure 3b, and 4 out of 15 & 17 flies for Figure 3c). We repeated this step for all possible combinations of test and training datasets, performing exhaustive leave p-out cross-validation, and calculated the fraction of correct classifications. As artifacts in region 35 lead to NaN values, we removed this region from this analysis.

### Response clustering

We concatenated the different stimulus response time series and calculated the average time series for each atlas region across animals. We then calculated the distance matrix based on Pearson correlations between time series and performed hierarchical clustering using Ward’s method. The order of atlas regions in Figure 2c is based on this clustering, responses within each region are ordered according to the average value. As artifacts in region 35 lead to `NaN` values, we removed this region from this analysis.

The response clustering in the correlation matrices in Figure 4e was performed as follows: in each protein deprived animal we calculated the correlations between all possible SEZ region pairs, based on the yeast stimulation time series. We then averaged the resulting correlation matrices across animals, and performed hierarchical clustering using Ward’s method. The correlation matrix for the fully fed state was calculated in the same way but ordered based on the clustering performed in the protein-deprived state.

### Response magnitude quantification

Responses were calculated as the average ΔF/F value during the 5 s of the first stimulation.

### Response onset

Response onsets in Figure 4f and g were defined as the time point after the first and before the second stimulation at which fluorescence first exceeded 4 × the SD of the background activity.

### Difference of mean responses between fed and protein deprived animals

Bootstrapping analysis in Figure 5c and e has been performed on response magnitude data by randomly sampling data points with replacement. We took 10000 bootstrap samples to calculate the mean and the confidence interval (bias-corrected and accelerated) of difference of means comparing fed to protein-deprived animals. We used the Python library `DABEST` (Ho et al., 2019) for performing the bootstrap analysis, and custom Python code using `Matplotlib` (Hunter, 2007) for the visualization.

### Volume renderings

SEZ volume renderings were created using the 3Dscript ImageJ plugin (Schmid et al., 2019).

### Bridge registration

To bridge the functional atlas to the JFRC2 standard, we performed recordings of the SEZ as well as the full central brain in the same animal, with our normal physiological and additional high-resolution imaging parameters. The physiological parameters were as described above, the high-resolution anatomical stack was recorded at an increased field of view and with increased z-resolution of 1 µm and with each plane imaged at 32 × plane averaging, resulting in a stack of 512 × 512 × 282 voxels at a resolution of ~0.5 × 0.5 × 1.0 µm. Using `ANTs`, we 1) aligned the SEZ scan (SEZp) to the SEZ standard, 2) SEZp to the central brain scanned with the physiological parameters (WBp), 3) WBp to the high-resolution anatomical scan of the central brain (WBa) and finally 4) we registered WBa to JFRC2. To improve registration, we added additional landmarks to the SEZ of JFRC2 by merging image data from 4 driver lines that selectively innervate the characteristic horn-like AMS1 regions. We obtained registered image stacks of the 4 lines (16B10, VT008676, VT044843, VT062247) from https://virtualflybrain.org. As our data consisted of *in-vivo* 2-photon scans of GCaMP6s baseline fluorescence whereas the JFRC2 standard consists of nc82 antibody stainings from fixated brains, we performed the last registration step with `ANTs` parameters optimized for cross-modal registration. Finally, we applied all resulting transforms to the functional atlas to transform it to JFRC2 space. Note that in the first step, we applied the inverse transform to move the functional atlas from SEZ standard to SEZp space. We validated the successful alignment by visually inspecting the overlap of neuropil borders including the antennal lobes, mushroom body lobes, fan-shaped body, ellipsoid body, as well as the AMS1 region and prominent neuronal tracts.

### Intersection with Braincode and driver line selection

To identify the Braincode regions with the largest overlap with the inner and outer borboleta regions, we downloaded the Braincode k60 clustering based on the Janelia FlyLight lines from https://strawlab.org/Braincode/. From the JFRC2 aligned version of our atlas we created image stacks containing only the inner and outer borboleta regions and downsampled this data to the dimensions of Braincode. In a custom-written Python script, we calculated the number of overlapping voxels between either borboleta regions and each Brain-code cluster. For the Braincode clusters with the highest overlap, we queried the Braincode database for driver lines with the highest coverage fraction. We then inspected expression patterns of these lines visually in the Janelia Fly-Light database and selected 8 candidate driver lines with reasonably sparse and specific expression in the inner or outer borboleta. Two lines that showed gross unspecific behavioral phenotypes (e.g. constant wing spreading) upon thermogenetic activation were excluded from the flyPAD analysis.

### flyPAD assays

flyPAD assays were performed based on a protocol previously described (Itskov et al., 2014) One well of the flyPAD was filled with 20 mM sucrose, and the other with 10 % yeast, each dissolved in a 1 % agarose gel. Flies were individually transferred to flyPAD arenas by mouth aspiration and allowed to feed for one hour at 25 °C (in Figure 1a and Figure S12a and b) or at 30 °C (in Figure 6 and Figures S11 and S12c and d), 70 % RH. flyPAD data were acquired using the Bonsai framework (Lopes et al., 2015), and analyzed in MATLAB using custom-written software, as described in (Itskov et al., 2014).

### Image-based tracking assay

Video tracking assays were performed based on a protocol previously described (Corrales-Carvajal et al., 2016). In brief: Flies were individually transferred to circular arenas by mouth aspiration and allowed to freely forage for one hour at 30 °C, 70 % RH while being video-recorded. The arena was equipped with 12 equally distributed food spots, six containing 10 % yeast and six 100 mM sucrose, each in 1 % agarose. Data were acquired using the Bonsai framework (Lopes et al., 2015), and analyzed in Python. Based on the validation in Corrales-Carvajal et al. (2016), we used micromovements as a proxy for feeding behavior. Micromovements are classified as segments, in which the fly’s head translational speed is below 2 mm/s and is closer than 2.5 mm to a given food spot. We excluded segments with translational speeds below 0.2 mm/s as resting, as well as segments with a duration shorter than a sip (0.1 s).

### Thermogenetic neuronal activation experiments

Flies carrying a transgene encoding the transient receptor potential channel *dTRPA1* gene under UAS control were crossed with candidate GAL4 lines in order to acutely activate distinct neuronal subpopulations in experimental flies. Flies were reared at 22 °C. 4 days after eclosion, female flies were sorted into fresh YBM and Canton-S males were added to ensure mating. Flies were kept at 22 °C on YBM for 10 days and transferred to fresh medium every 3 days. Activation was induced by preincubating flies at 30 °C, 70 % RH for two hours. flyPAD assays or the video tracking assays were then performed at this restrictive temperature for one hour.

### Immunohistochemistry and image acquisition

Males from each Gal4 line were crossed to females homozygous for the *UAS-CD8::GFP* reporter line and 3–10 day-old adult females carrying both the GAL4 driver and the UAS reporter were dissected. Samples were dissected in 4 °C PBS after a quick passage through EtOH, and were then transferred to formaldehyde solution (4 % paraformaldehyde in PBS + 10 % Triton-X) and incubated for 20–30 min at RT. Samples were then washed three times in PBST (0.5 % TritonX in PBS and then blocked in Normal Goat Serum 10 % in PBST for 15-60 min at RT. Samples were then incubated in primary antibody solutions (Rabbit anti-GFP [Torrey Pines Biolabs, Secaucus, NJ, USA] at 1:6000 and Mouse anti-NC82 [Developmental Studies Hybridoma Bank] at 1:10 in 5 % Normal Goat Serum in PBST). Primary antibody incubations were performed for 3 days at 4 °C with rocking. They were then washed in PBST 2–3 times for 10–15 min at RT. The secondary antibodies were applied (Anti-mouse A594 [Invitrogen] at 1:500 and Anti-rabbit A488 [Invitrogen] at 1:500 in 5 % Normal Goat Serum in PBST) and brains were then incubated for 3 days at 4 °C. They were again washed in PBST 2–3 times for 10–15 min at RT. Samples were mounted in Vectashield (Vector Laboratories, Burlingame, CA, USA). Images were captured on a Zeiss LSM 710 using a 25× glycerol objective.

## Author Contributions

D.M. and C.R. conceived and designed the project, interpreted data, and wrote the manuscript. D.M. performed experiments and data analysis. D.G. performed and analyzed video tracking experiments and performed bootstrapping analysis.

