## Supplementary Figures for "Distinct internal states interact to shape food choice by modulating sensorimotor processing at global and local scales"

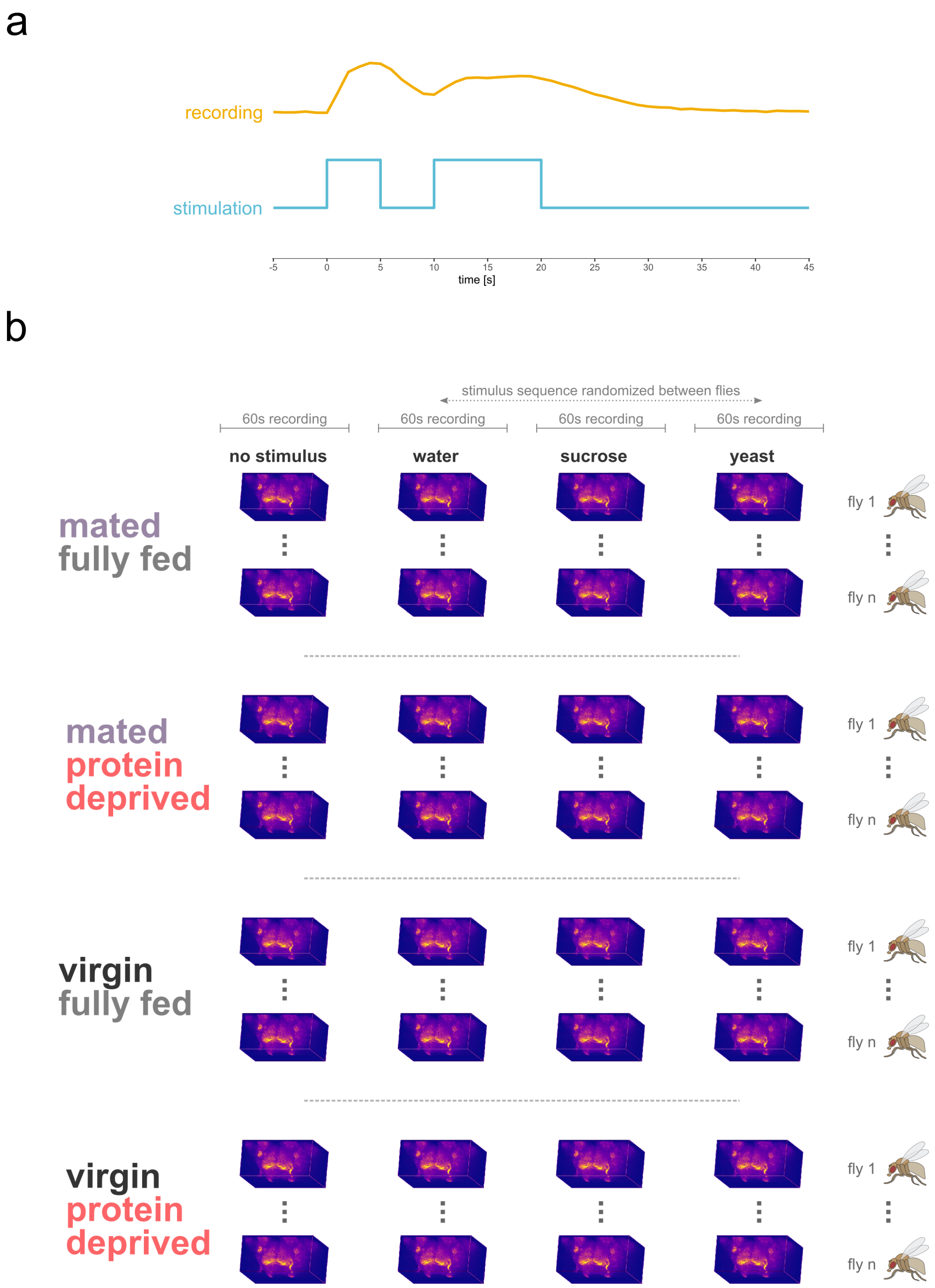


**Figure S1. Experimental design.** **a**, Per recording two consecutive taste stimulations lasting 5 s and 10 s respectively were applied to the proboscis. **b**, Experiments were performed in flies in four different internal state combinations. Experiments started with a recording without any stimulation and were followed by stimulations with three different taste stimuli (water, sucrose & yeast) that were applied in randomized order.

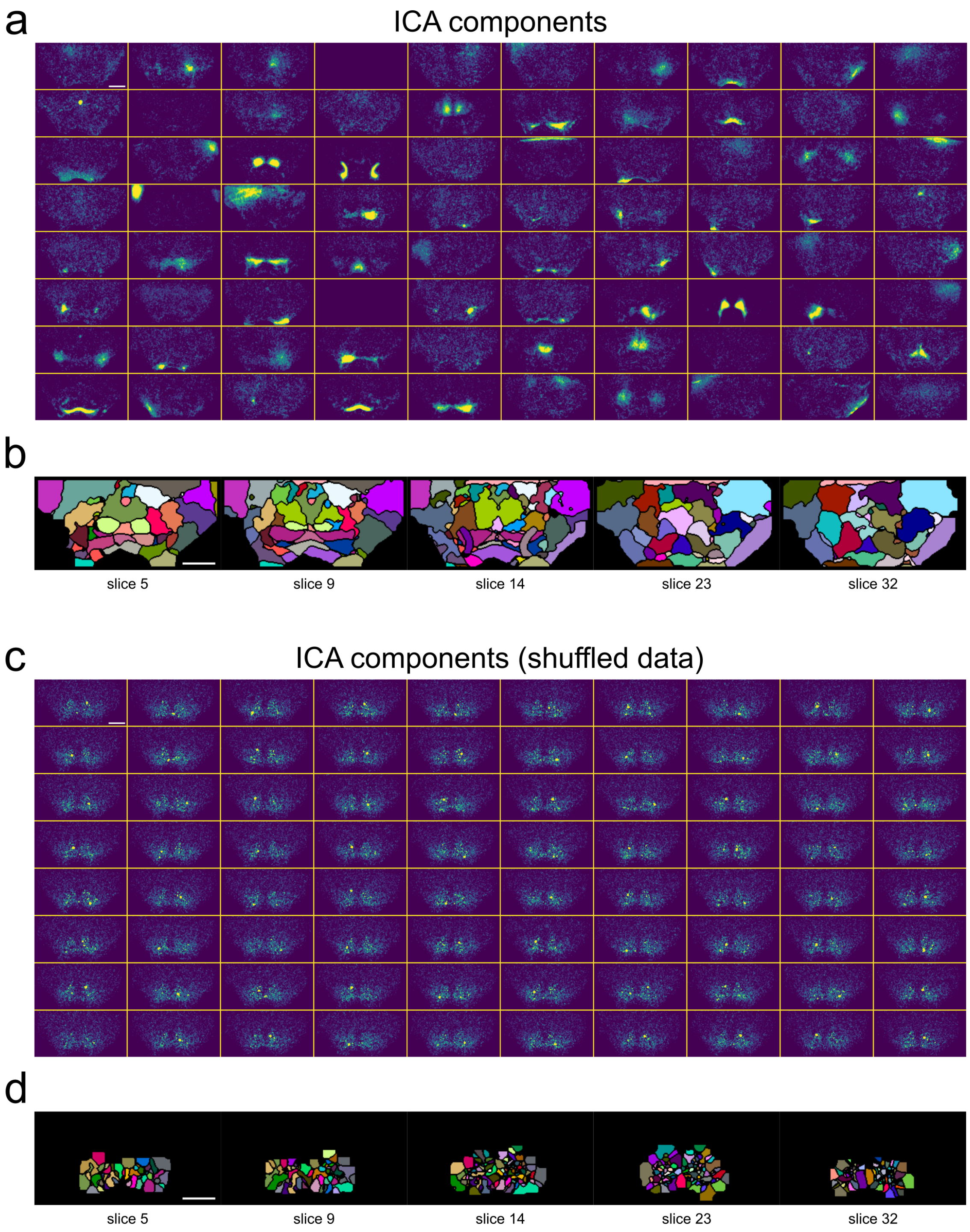


**Figure S2. Extracted independent components.** **a**, Maximum intensity projections of the 80 independent components identified by the spatial ICA algorithm (Varoquaux et al., 2010). **b** Consecutive sections of the resulting binary SEZ atlas after performing the atlas building steps. For details see Material and Methods. **c**, Maximum intensity projections of the resulting independent components after running the ICA on temporally shuffled data using the exact same settings as in **a**. **d**, as in **b** but atlas building performed on data plotted in **c**. Scale bars correspond to 50 µm.

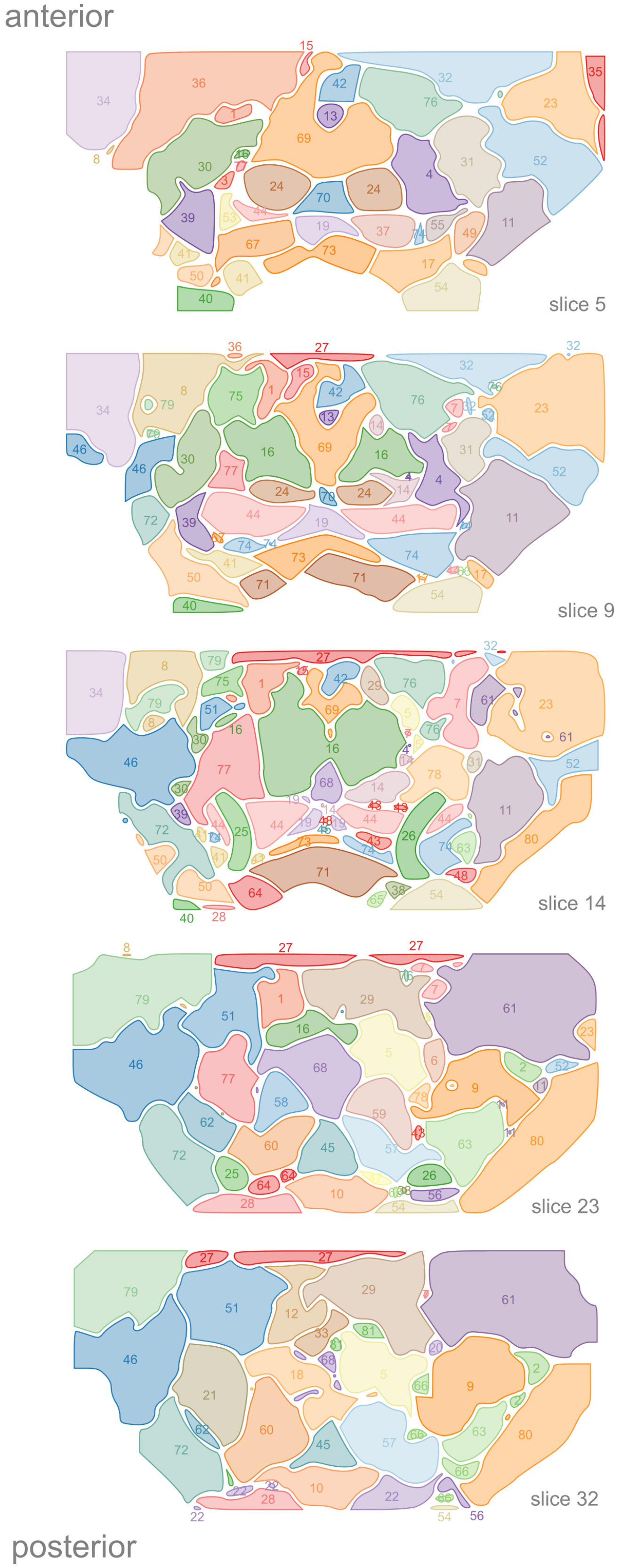


**Figure S3. Region IDs of the functional SEZ atlas.** Five slices of the functional SEZ atlas ordered from anterior (top) to posterior (bottom), indicating the corresponding atlas region numbers.

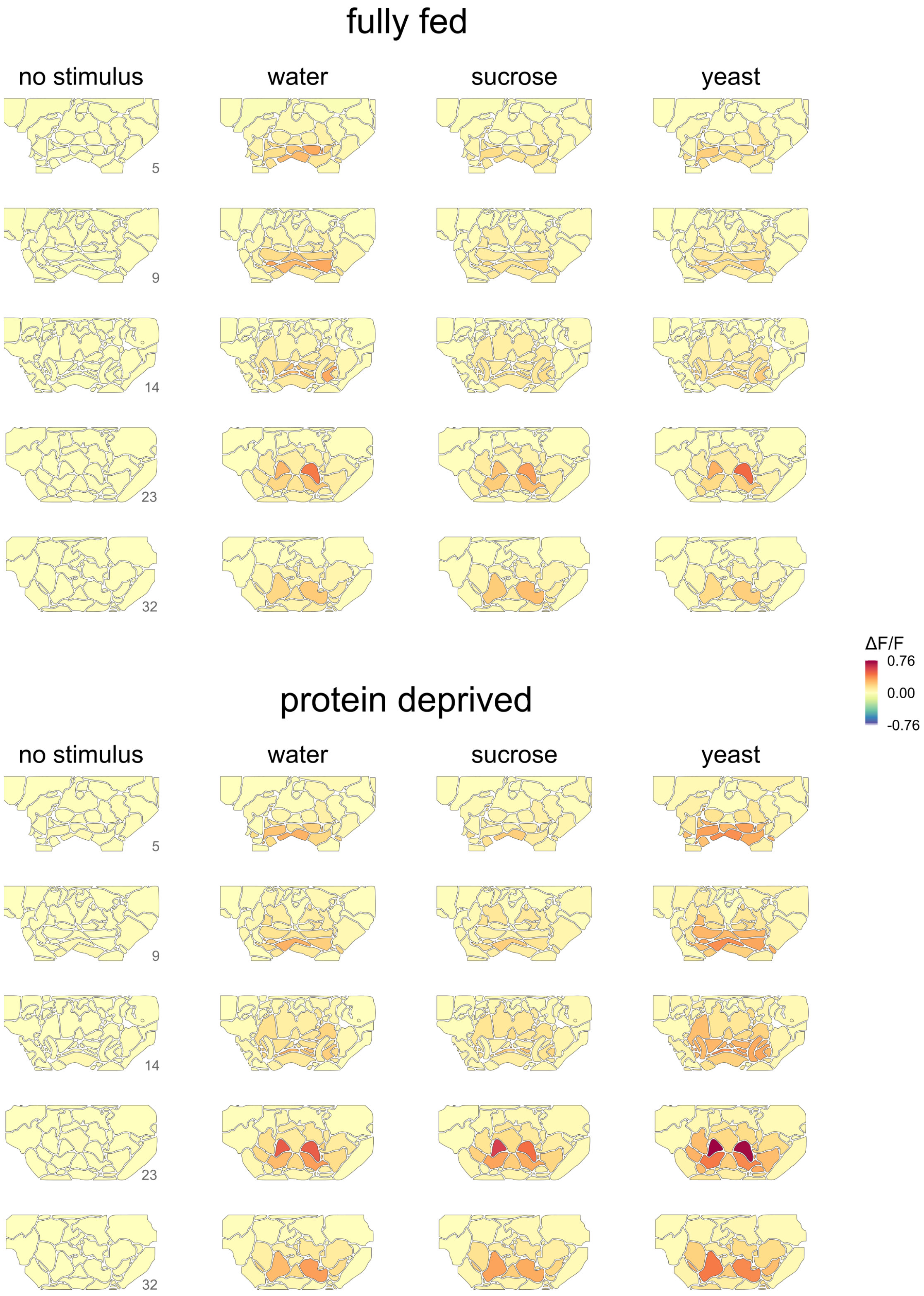


**Figure S4. Stimulus response maps at different metabolic states.** Stimulus-response maps for no stimulus and three different stimulations. Colors indicate median ΔF/F response values across animals during the first stimulation. Numbers indicate slice numbers. n = 7-10.

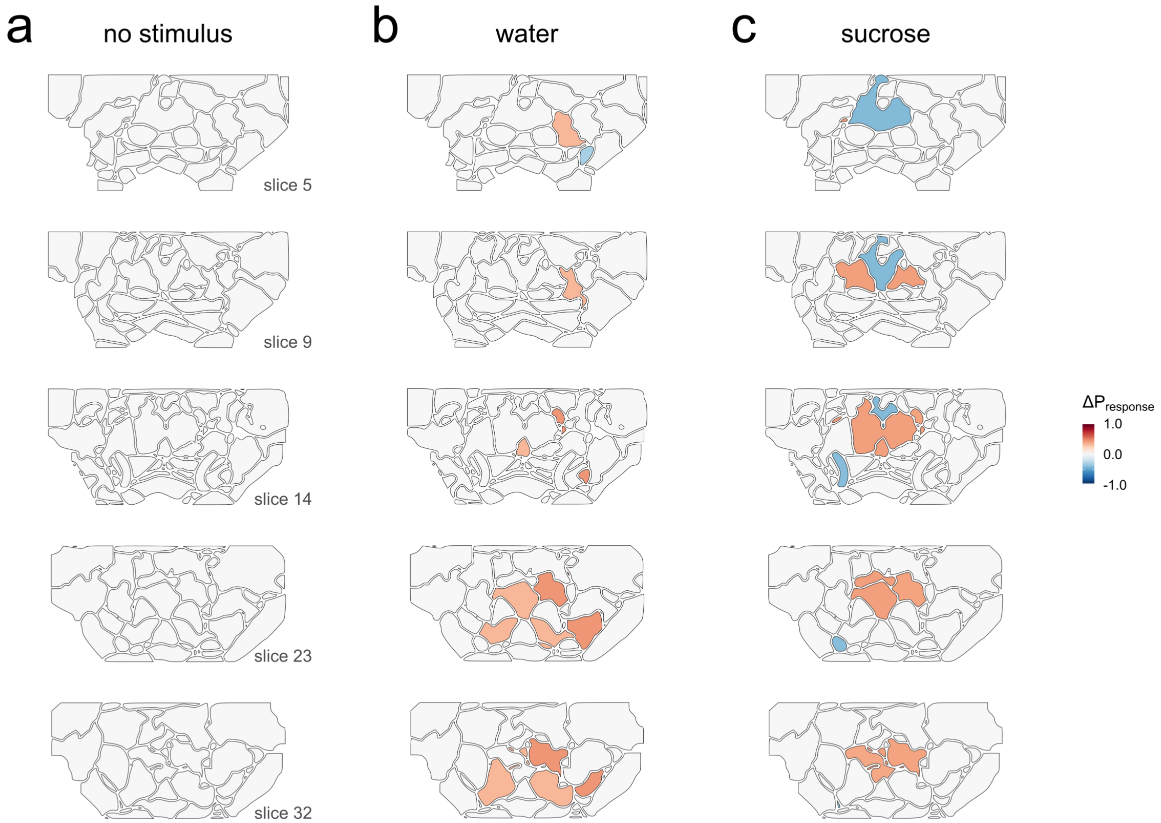


**Figure S5. Changes in response probability for different taste stimuli.** The change in response probability across SEZ regions upon protein deprivation for different food taste stimuli, mapped onto the SEZ atlas. Only changes in response probability that exceeded ± 0.3 are plotted.

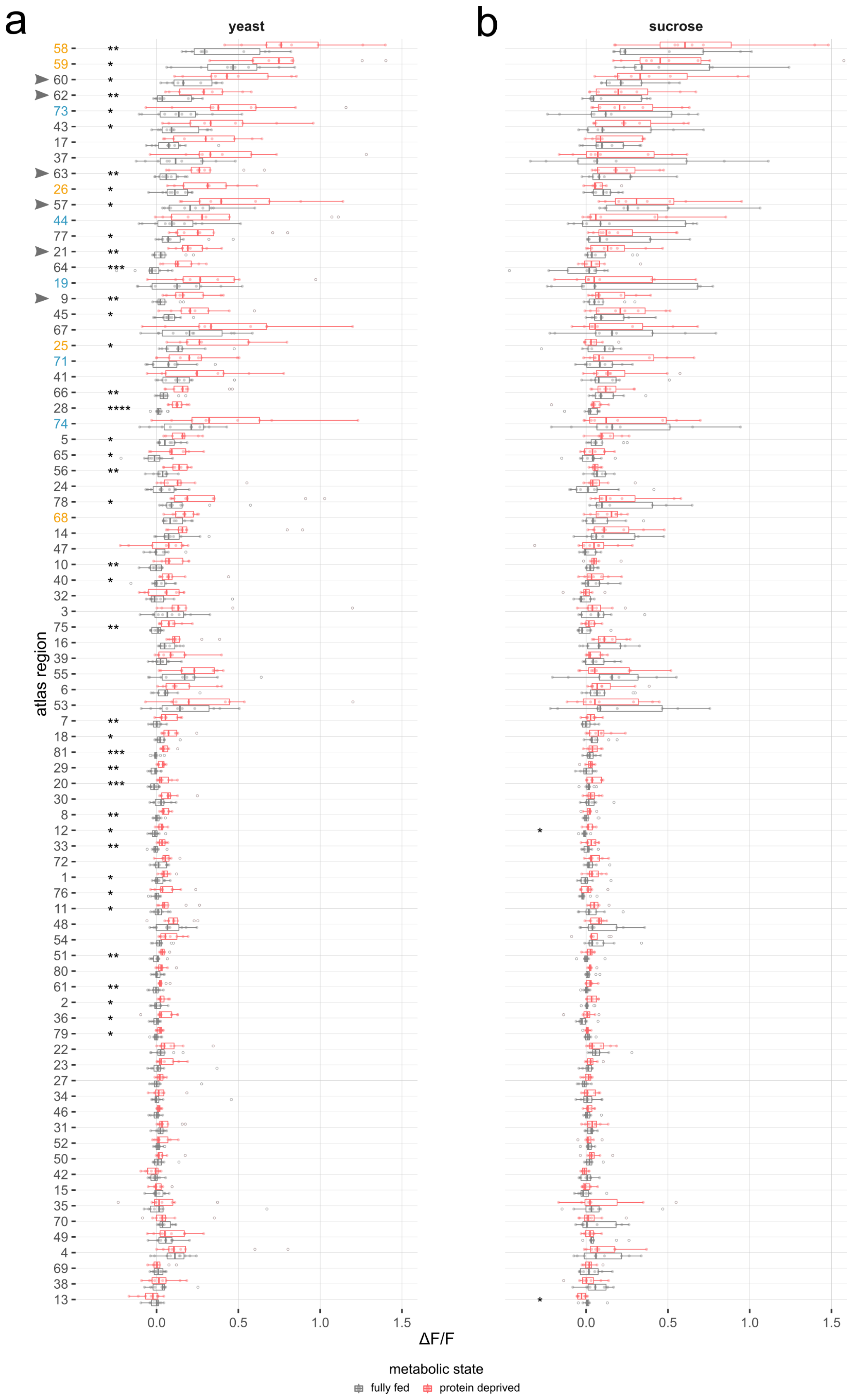


**Figure S6. Response modulation for all atlas regions by metabolic state.** Boxplots of yeast (**a**) and sucrose (**b**) stimulus-induced responses across all atlas regions, compared between metabolic states in mated females. The data is ordered according to the difference of median values between corresponding fully fed (*gray*) and protein-deprived (*red*) flies stimulated with yeast. *yellow* = sensory regions, *blue* = motor regions, *arrowheads* indicate borboleta regions. Points indicate ΔF/F response values in different experiments. Boxplots indicate median, first, and third quartile, whiskers extend to the lowest and highest values that lie within 1.5 of the inter-quartile range of the box. Wilcoxon rank-sum test, n = 7-10, * $P < 0.05$, ** $P < 0.01$, *** $P < 0.001$, **** $P < 0.0001$. Scale bars correspond to 50 µm.

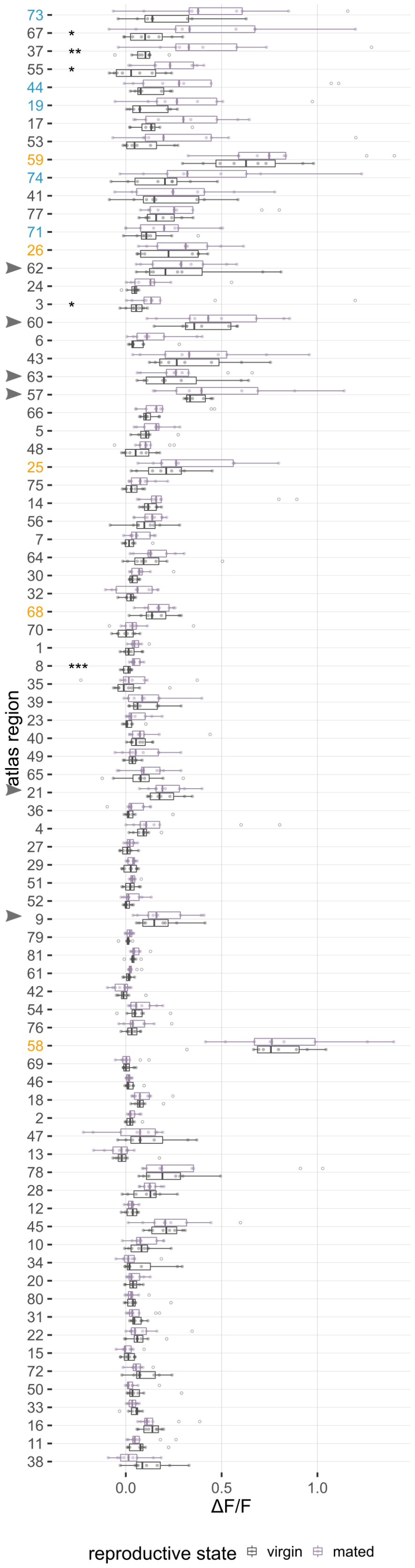


**Figure S7. Response modulation for all atlas regions by reproductive state.** Boxplots of yeast stimulus-induced responses across all atlas regions, compared between reproductive states in protein-deprived females. The data is ordered according to the difference of median values between corresponding virgin & protein-deprived flies (*gray*) and mated & protein-deprived flies (*purple*). *yellow* = sensory regions, *blue* = motor regions, *arrowheads* indicate borboleta regions. Points indicate ΔF/F response values in different experiments. Boxplots indicate median, first, and third quartile, whiskers extend to the lowest and highest values that lie within 1.5 of the inter-quartile range of the box. Wilcoxon rank-sum test, n = 8-9, * $P < 0.05$, ** $P < 0.01$, *** $P < 0.001$.

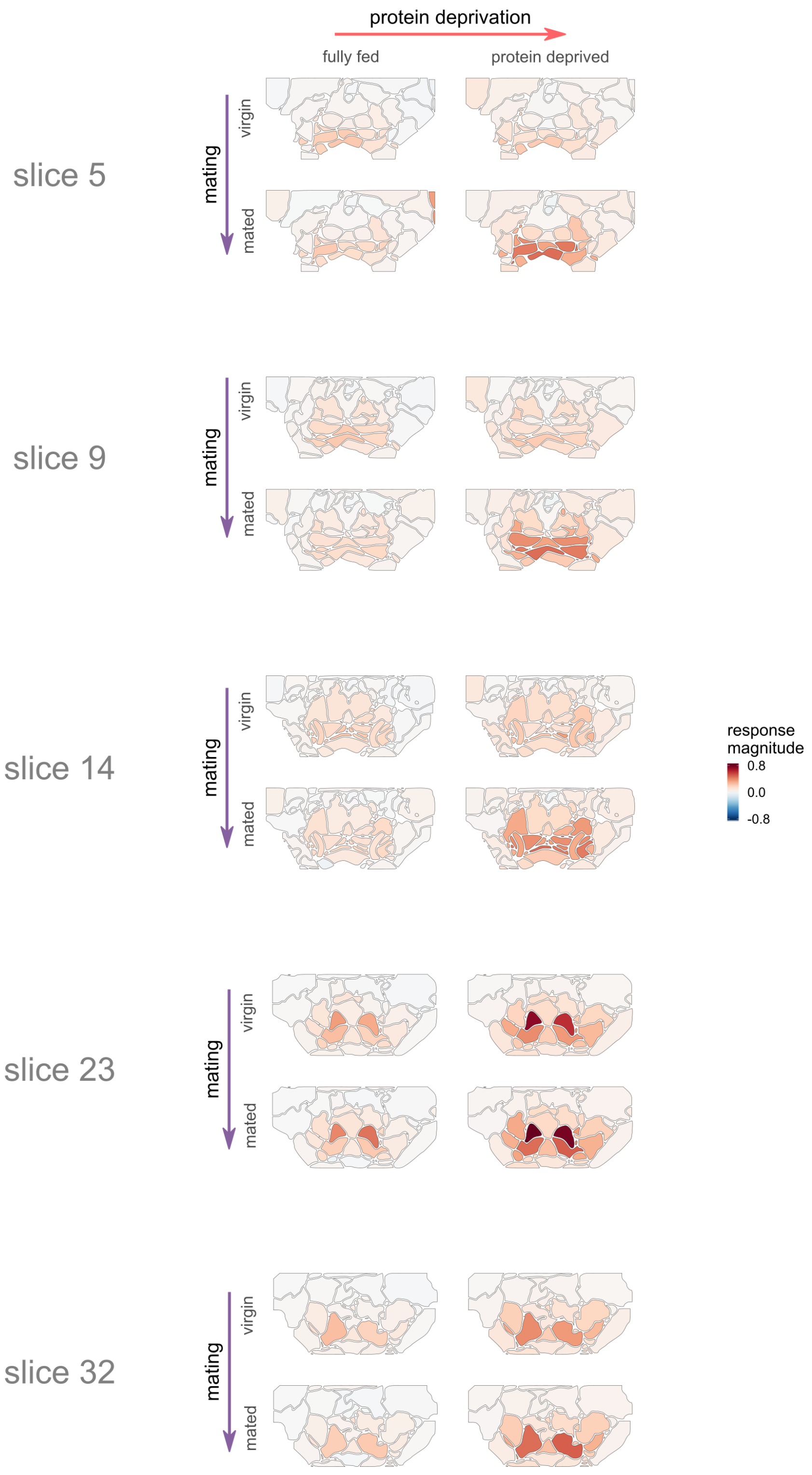


**Figure S8. Yeast stimulus response maps for four internal states.** Stimulus-response maps for yeast stimulations in four internal states (fully fed & virgin, fully fed & mated, protein deprived & virgin, protein deprived & mated). Colors indicate mean ΔF/F response values across animals during the first stimulation. n = 7-10.

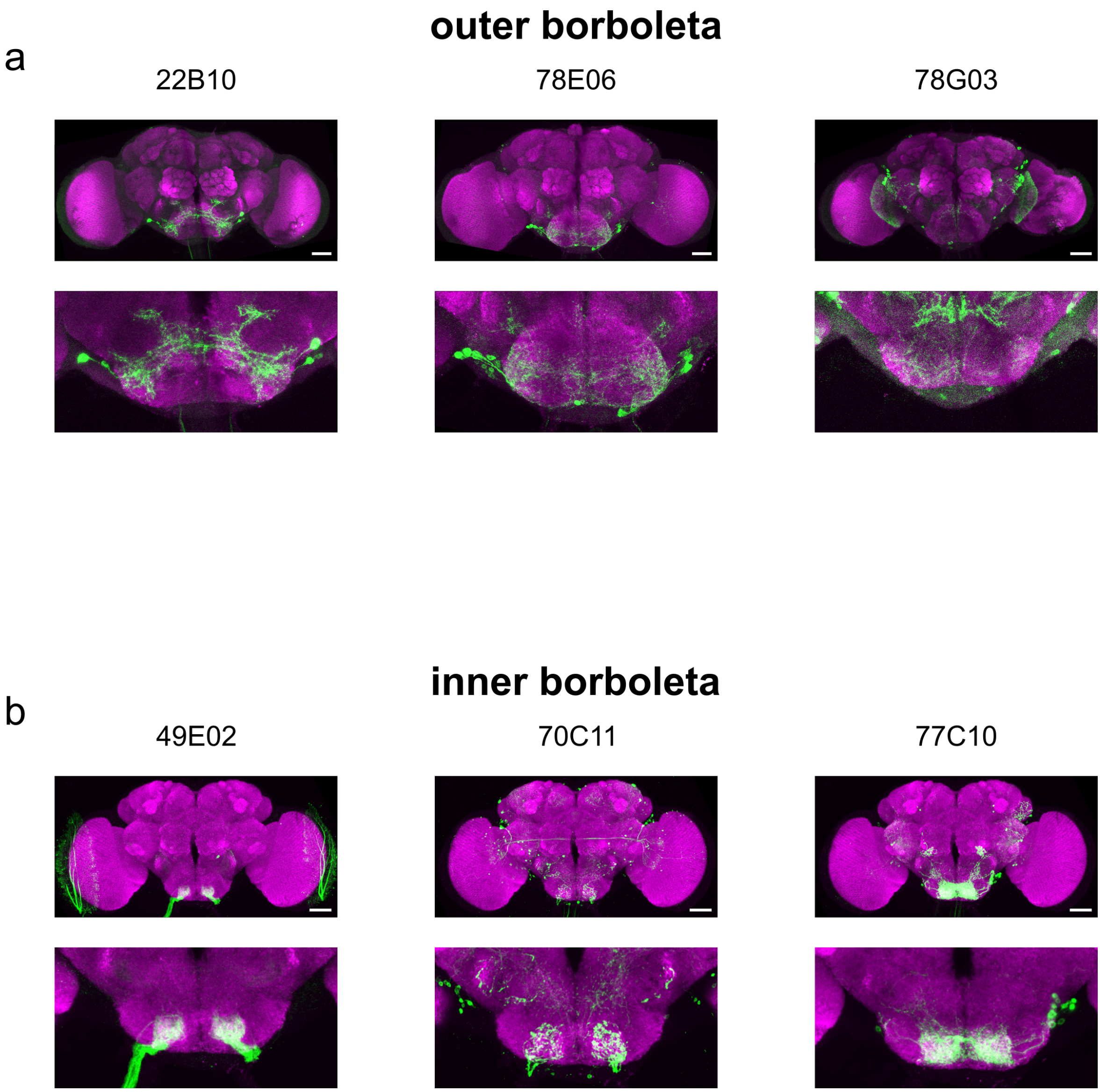


**Figure S9. Anatomy of borboleta-innervating Gal4 lines.** **a**, Anatomy of three Gal4 lines that innervate the outer borboleta region. *top:* maximum intensity projections of full confocal brain scans, *bottom:* zoom into the SEZ region. **b**, Anatomy of three Gal4 lines that innervate the inner borboleta region. *top:* maximum intensity projections of image stacks obtained from virtualflybrain.org, *bottom:* zoom into the SEZ region. Scale bars correspond to 50 µm. Gal4 expression pattern (*UAS-CD8::GFP*) is shown in *green*, nc82 neuropil staining is shown in *magenta*.

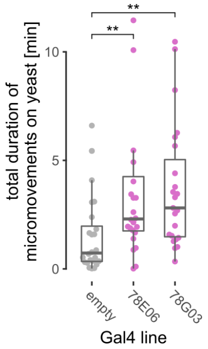


**Figure S10. Video tracking experiments of borboleta activation.** Total duration of micromovements on yeast and sucrose spots as measured by image-based tracking of mated female fully fed flies maintained for 2 h at restrictive temperature. Points indicate the total duration of micromovements flies performed on yeast food spots. Boxplots indicate median, first, and third quartile, whiskers extend to the lowest and highest values that lie within 1.5 of the inter-quartile range of the box. Wilcoxon rank-sum test, $n = 21\text{-}24$, ** $P < 0.01$. Some outliers were excluded from plotting due to scaling.

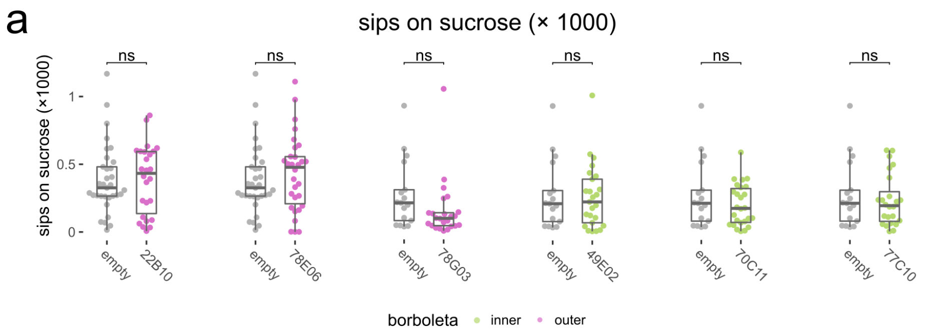


**Figure S11. Sucrose feeding effects observed in borboleta activation experiments.** Number of sips from 20 mM sucrose as measured by the flyPAD assay by mated female fully fed flies maintained for 2h at restrictive temperature. These data correspond to the experiments shown in Figure 6. The genotype of the Gal4 lines used to drive TRPA1 expression is depicted below the plots with empty depicting the matching empty-Gal4 control. Outer borboleta lines in magenta and inner borboleta lines in green. Points indicate the average number of sips on sucrose for each fly. Boxplots indicate median, first, and third quartile, whiskers extend to the lowest and highest values that lie within 1.5 of the inter-quartile range of the box. Wilcoxon rank-sum test, $n = 17$-$33$, ns $P > 0.05$.

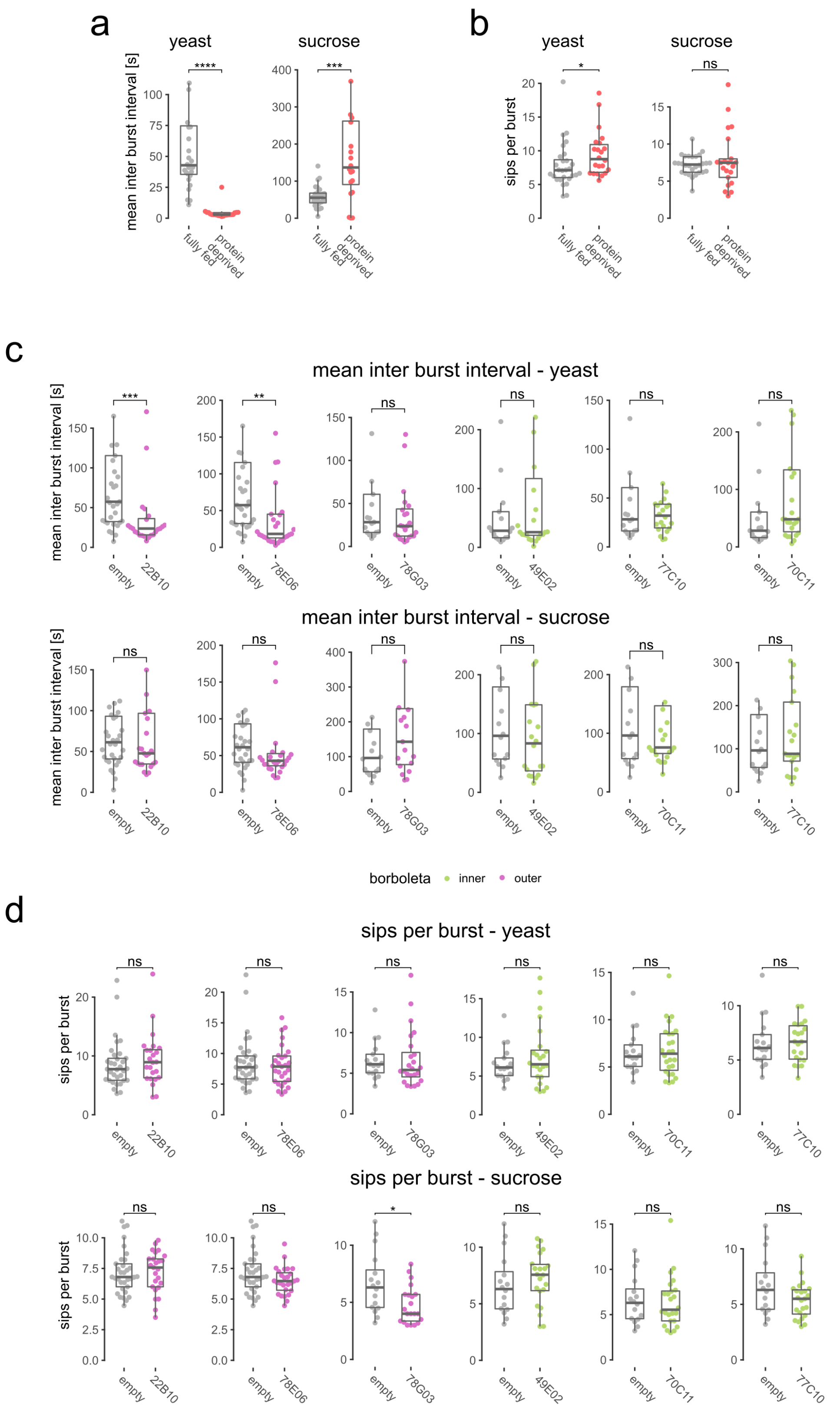


**Figure S12. Feeding microstructure effects observed in borboleta activation experiments. a** and **b**, Feeding microstructure parameters for 57C10-Gal4,UAS-GCaMP6s flies as measured using the flyPAD assay. Points indicate the mean inter-burst interval in seconds (**a**) and sips per feeding burst (**b**). Boxplots indicate median, first, and third quartile, whiskers extend to the lowest and highest values that lie within 1.5 × the inter-quartile range from the box. Wilcoxon rank-sum test, n = 20-28, * $P < 0.05$, ** $P < 0.01$, *** $P < 0.001$. ns $P > 0.05$. **c**, Mean inter-burst interval as measured by the flyPAD assay from 10 % yeast (top) and 20 mM sucrose (bottom), by mated female fully fed flies maintained for 2h at restrictive temperature. The genotype of the Gal4 lines used to drive TRPA1 expression is depicted below the plots with empty depicting the matching empty-Gal4 control. Results for lines showing expression in the outer borboleta are plotted in *magenta* and for lines showing expression in the inner borboleta are plotted in *green*. Points indicate the mean inter-burst interval in seconds for each fly. Wilcoxon rank-sum test, n = 16-33. **d**, As **c** but for the number of sips per burst. All plots correspond to the analysis of the experiments shown in Figure 6. Wilcoxon rank-sum test, n = 16-33. Some outliers were excluded from plotting due to scaling.

**Data S1. SEZ atlas** 3D NifTI image stack of the binary SEZ atlas used for signal extraction.

**Data S2. SEZ standard** 3D NifTI image stack of the SEZ standard used for alignments.

**Video S1. Volume rendering of volumetric imaging data** Example recording showing the SEZ response upon yeast stimulation in a protein deprived fly. The recording is looped and the z-plane is changed from anterior to posterior to show responses across different SEZ layers. The stimulation period is indicated in the upper left corner. Playback speed is 10×.
